# Antibiotic-Specific Conformational Landscapes of a Multidrug Transporter

**DOI:** 10.1101/2025.05.17.654648

**Authors:** Hassan Maklad, Tom Kache, Aurélie Roth, Maria Mamkaeva, Cédric Govaerts, Jelle Hendrix, Chloé Martens

## Abstract

Multidrug transporters are membrane proteins that can transport an ensemble of structurally dissimilar compounds and contribute to bacterial multidrug resistance (MDR) by exporting different antibiotics from the cell. However, whether they transport different substrates through a common mechanism or via distinct substrate-dependent mechanisms remains unclear. In this work, we used single-molecule Förster resonance energy transfer (smFRET) to measure time-resolved conformational dynamics of LmrP, a multidrug transporter of the Major Facilitator Superfamily (MFS). We present high-resolution conformational landscapes of LmrP in the presence of different antibiotics. Through multi-parameter Hidden Markov Modeling (mpH^2^MM), we uncovered transient states and quantified their sub-millisecond interconversion kinetics. We observed antibiotic-dependent heterogeneity in the conformational landscape, both in accessible states and in interconversion rates. Notably, poorly or non-transported antibiotics slow down transition kinetics, pointing to rapid state interconversion as a driver of efficient transport. This suggests that MFS MDR transporters bind and export structurally dissimilar antibiotics by relying on an array of underlying conformational states with ligand-dictated interconversion rates. This work provides novel insights into the mechanism of MDR transporters and advocates for combined structure/dynamics-based drug design when targeting their function.

## Introduction

Active extrusion of drugs across the cell membrane via transport proteins is a key driver of multidrug resistance in bacterial pathogens^1–3^. By decreasing the intracellular drug concentration, multidrug transporters enable bacteria to survive upon antibiotics treatment and eventually acquire genetically encoded resistance mechanisms, such as target mutation or enzymatic degradation. A molecular-level understanding of the transport mechanism of efflux pumps is a prerequisite for the development of targeted inhibition strategies to re-sensitize resistant strains ^4,5^.

How a single transport protein can recognize and translocate substrates with widely diverse chemical properties such as charge, size, and shape remains a fascinating enigma and, at the same time, complicates the rational design of efflux pump inhibitors. Structural adaptation to chemically diverse substrates is observed for the prototypical efflux pump AcrB, which features distinct binding pockets and multiple substrate pathways^6–8^. AcrB and its homologs, which belong to the same structural family of Resistance-Nodulation-Division (RND) transporters, play a prominent role in the emergence of multidrug-resistant phenotypes in Gram-negative bacteria ^9,10^. These tripartite pumps are absent in Gram-positive bacteria, where most drug efflux systems belong to the Major Facilitator Superfamily (MFS)^1–3,11,12^.

The molecular architecture of MFS efflux pumps differs markedly from that of RND pumps. MFS transporters are composed of 12 to 14 transmembrane helical segments arranged in two bundles of six helices that surround a central binding pocket ^4,13,14^. MFS proteins mediate transport via the alternating access model, where the transporter alternates between inward- and outward-facing states to sequentially expose the binding site to each side of the membrane ^15^. How these multidrug efflux pumps reconcile broad ligand promiscuity with a conserved MFS fold is poorly understood: do different substrates follow distinct pathways through conformational space, or do they converge onto a common sequence of structural rearrangements leading to transport? As a corollary, how multidrug transporters distinguish between transported and non-transported binders remains unclear.

Here, we address these questions using the multidrug transporter LmrP from *Lactococcus lactis*^16^. LmrP is a substrate/proton antiporter from the Major Facilitator Superfamily that can export several molecules including antibiotics^17,18^. We have previously investigated the coupling of LmrP conformational changes to protons and ligand binding using Double Electron-Electron Resonance (DEER) distance measurements on paramagnetically labelled protein mutants ^19–21^. These studies have shown that LmrP alternates between at least two conformational states, an inward-open (IO) and an outward-open (OO) state, where the protonation states of key acidic residues orchestrate the transition^19^ such as Asp68, a highly conserved residue in the MFS family^22,23^. n parallel, X-ray crystallography characterization and DEER studies suggest that LmrP binds different ligands in an OO conformation, consistent with a “one-size-fits-all” high affinity state^24^. These data were obtained from solid (frozen) samples and thus conformationally fixed protein samples, thereby excluding the range of dynamically interconverting structural states, including short-lived ones, that compose the conformational landscape of the transporter. Recent studies on transporters have revealed that heterogeneous conformational landscapes composed of multiple subpopulations define transport function^25–28^. Whether this underlying complexity can be leveraged to accommodate multidrug binding and, subsequently, multidrug transport remains an open question.

Practically, an accurate depiction of a conformational landscape requires identifying pathways between different conformations occupied for different time periods. To identify the different states, the pathways that connect them and the underlying kinetics of the transitions, we turned to single-molecule Förster resonance energy transfer (smFRET) which is uniquely positioned in that regard^29–31^. In FRET, a non-radiative energy transfer occurs from a donor fluorophore (D) to an acceptor fluorophore (A), where both are attached to specific sites on a protein molecule ^28,32^. The efficiency of the energy transfer, *E*, sensitively reports on the distance between the donor and acceptor on the nanometer length scale^33^. Real-time tracking of *E* for single molecules in turn allows monitoring protein conformational changes and dynamics at nanometric spatial resolution combined with a wide range of accessible timescales (ns-minutes).^34^ Specifically, we employed multi-parameter fluorescence detection with Pulsed-Interleaved Excitation characterization (PIE-smFRET)^35^(**Fig.1A**) to study the conformational dynamics of LmrP and characterize its ligand-dependent conformational landscape. Our work reveals how distinct conformational states and their interconversion kinetics together form a fine-grained conformational landscape that different ligands navigate in distinct ways.

**Figure 1.**
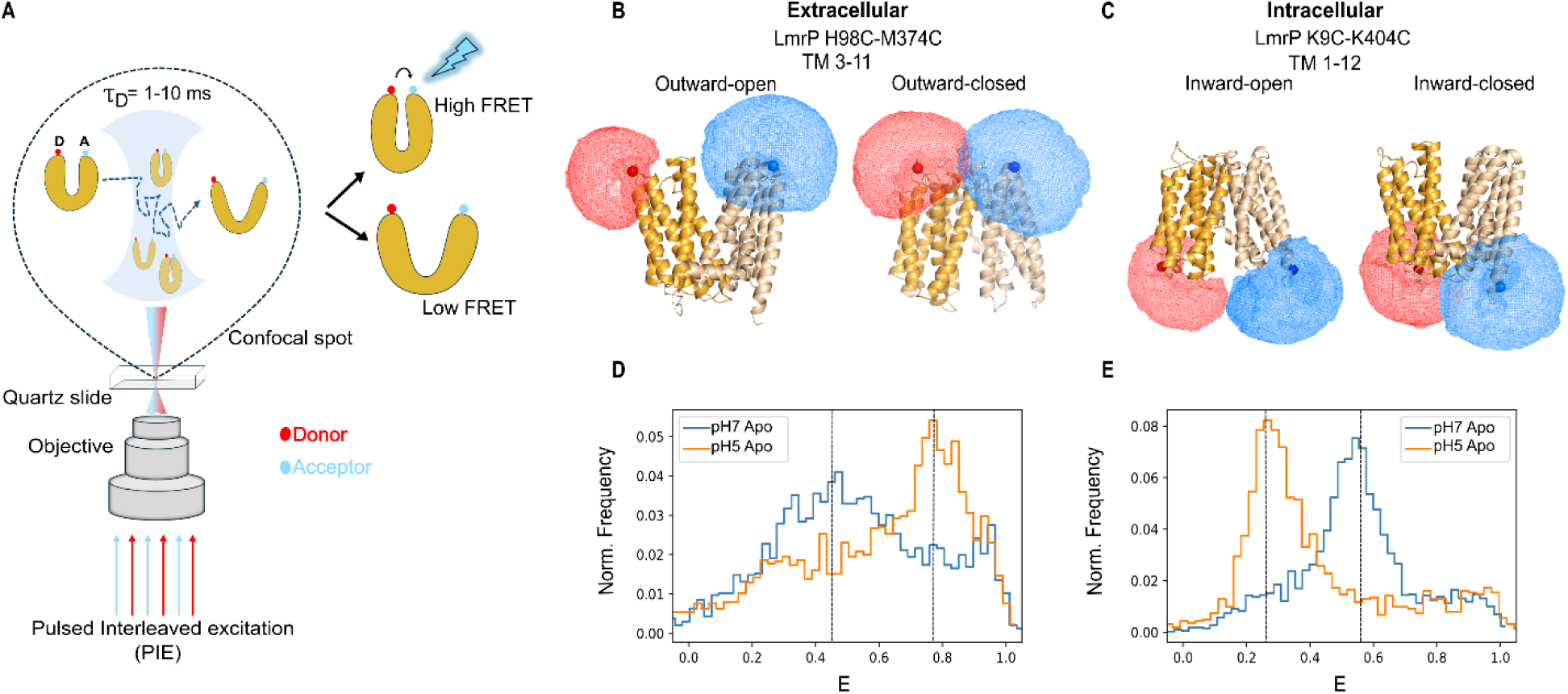
smFRET charts the conformational landscape of LmrP. **A.** Schematic illustration of the PIE smFRET experiment setup, showing the free diffusion of labelled LmrP molecules across the focal spot of an inverted fluorescence microscope. “D” and “A” correspond to donor and acceptor dyes, respectively. In the PIE setup, two spatially overlapped and alternatingly pulsing lasers are shooting at the objective to excite donor and acceptor fluorescence consecutively. Fluorescence signals of the donor and acceptor are recorded separately, and the fluorescence lifetime and anisotropy of each dye are determined.**B and C.** Cartoon representation of LmrP helices with 98C (TM3)-374C (TM11) in outward-open **(OO)** or outward-closed **(OC)** conformations, and LmrP 9C (TM1)-404C(TM12) in inward-open **(IO)** or inward-closed **(IC)** conformations. The accessible volumes of the fluorophores ATTO488 (donor, red) and Alexa647 (acceptor, blue) are depicted as spheres. The N- and C-terminal halves of LmrP comprise TM1–6 (light orange) and TM7–12 (wheat beige), respectively. **D and E.** Burst-wise FRET efficiency histograms showing the smFRET signatures of intracellular and extracellular distance reporter mutants. The data reveals pH-dependent conformational changes of apo LmrP, with distributions recorded at **pH 7 (blue)** and **pH 5 (orange)**.

## Results

### Charting the conformational landscape of LmrP

Based on the crystal structure of LmrP in the OO conformation and the AlphaFold2 model in the IO conformation^36^, we selected pairs of residues with a spatial separation allowing to distinguish the two states by smFRET when using ATTO 488 (donor) and Alexa Fluor 647 (acceptor) dyes. Specifically, one pair was selected on the extracellular side (H98 and M374 in TM3 and TM11 respectively) and one at the intracellular side (K9 and K404 in TM1 and TM12) (**Fig.1B,C**). The selected residues were replaced with cysteines into a cysteine-less LmrP to enable covalent attachment of the fluorescent dyes through their maleimide groups. The predicted FRET efficiency values *E*, together with the corresponding expected distances between the probes in both conformations, are given in (**Table S.1**). All double cysteine mutants were functionally tested by following LmrP-mediated transport of Hoechst33342 in inverted membrane vesicles (**Fig. S1**). Single-molecule FRET measurements were performed on the labeled LmrP molecules in detergent micelles, diluted to ∼50-pM concentration, and freely diffusing through the femtolitre-size probe volume of the confocal microscope (**Fig.1A**). The distribution of FRET efficiencies for all measured molecules was plotted as burst-wise *E* histograms.

At pH 7, the conformational landscape of apo-LmrP is characterized by a broad distribution of *E* on the extracellular side, with a maximum around *E* ≈0.45 (**Fig. 1D, left, blue histogram**). On the intracellular side, the histogram displays a main central state with a maximum around *E*≈0.55 (**Fig.1E**, **right, blue histogram**). By comparing the observed to expected *E* values (**Table S.1**), we can conclude that the apo protein at pH 7 adopts a flexible conformational state that is predominantly closed on the intracellular side, while sampling different opening magnitudes on the extracellular side.

As changes in pH modulates the conformational equilibrium of LmrP^19^, we then performed smFRET measurements at pH 5, where we observe a shift in the distance distributions compared to pH 7. On the extracellular side the population centers around *E* ≈0.8, indicating closure upon lowering the pH (**Fig. 1D, left, orange histogram**) while, on the intracellular side, we observe the main population shifting to *E*≈0.3, indicating opening (**Fig. 1E, right, orange histogram).** We confirmed this observation by coupling the extracellular reporter with the D68N mutation, which is known to stabilize LmrP in the IO state ^19^ showing an increase of the high-FRET population at both pH 7 and 5, compared to wild-type (**Fig. S2)**. These observations are in line with our previous DEER studies^19,20^ and validate the ability of smFRET to identify and distinguish different conformational states of LmrP. Importantly, smFRET analyses up to this point have reported burst-averaged values of *E*. However, if molecules interconvert rapidly between two (or more) FRET states while diffusing through the probe volume, these states appear as averaged histograms rather than distinct peaks, leaving conformational intermediates with potential functional significance unresolved^37,38^.

### Kinetic analysis of apo LmrP reveals transient hidden states

To characterize the conformational states underlying the burst-averaged FRET histograms and the time scales at which these states interconvert, we took advantage of the advanced capabilities of the optical setup used in this work (*see Methods*). The setup records individual photons emitted by the donor and acceptor fluorophores, after their respective pulsed excitation, on the level of individual single-molecule fluorescence bursts. This enables the use of multiparameter photon-by-photon hidden Markov modeling (mpH^2^MM) ^35,39,40^ to monitor FRET changes in these bursts with sub-millisecond precision ^39^ (**Fig. S3**). Specifically, mpH^2^MM infers the number of conformational states a single molecule visits within a single burst, their mean *E* values, the stoichiometry *S*, the kinetic rate constants that report on how often molecules transition between given states and an estimated mean dwell time that corresponds to the average time that a molecule stays in a particular state before transitioning to another state. By integrating state identity and dwell times, we obtain a high-resolution conformational landscape for both the intracellular and extracellular reporters.

At pH 7, a three-state model was the best fit to describe the data for the distance reporters on either side of the membrane (**Fig. 2A**). When comparing the observed mean E values for each of the three states to those predicted from structural models (**Supplementary Table 1),** we observe only partial overlap. This discrepancy indicates that the protein in solution samples other states than the OO observed by crystallography or the IO predicted by Alphafold 2 modelling. We here annotated the three FRET states according to their mean *E* value: a low-FRET state **L** (*E* = 0.1–0.4), a medium-FRET state **M** (*E* = 0.4–0.7), and a high-FRET state **H** (*E* = 0.7–1.0). A subscript (E or I) is used to specify whether the measurement is on the extracellular or intracellular reporter.

**Figure 2.**
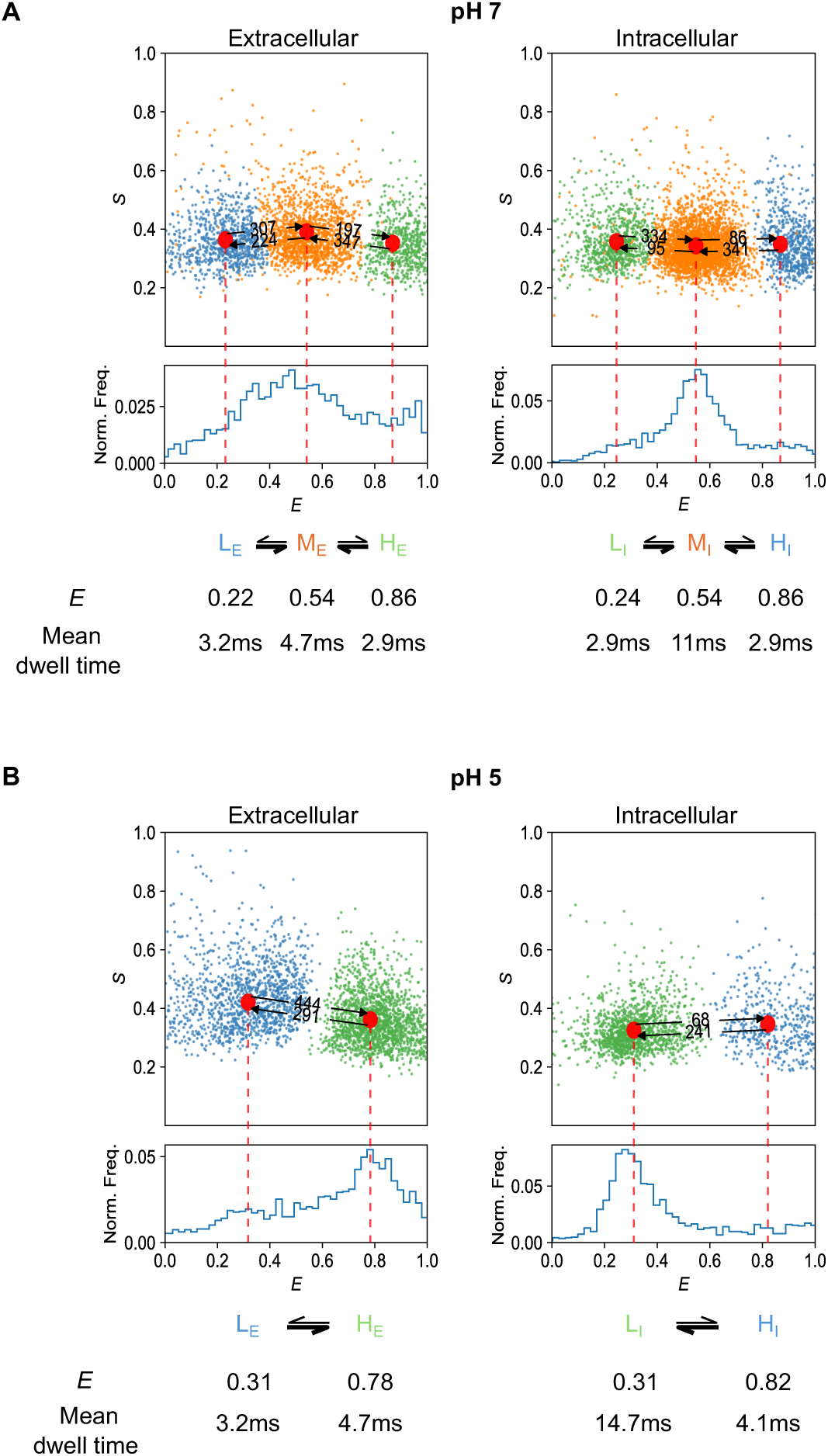
Kinetic analysis of apo LmrP reveals transient hidden states. **A.** mpH²MM analysis of LmrP in the apo state at pH 7. *Upper panel:* E–S scatter plot of individual dwells, colored according to the state assignment of the selected mpH²MM model. Red circles indicate the mean FRET efficiency (E) of each state, and the numbers above the arrows denote the transition rate constants (s⁻¹) between FRET states. *Lower panel:* Burst-wise FRET histogram showing corrected, exclusively FRET-active bursts (see Methods). The mean E value and mean dwell time (ms) for each interconverting FRET state are indicated below. Data from extracellular distance reporters are shown on the left, and intracellular distance reporter data on the right. **B**. mpH²MM analysis of LmrP at pH 5. Data are presented using the same layout and analysis as in panel **A**.

We observe that the interconversion kinetics between the three states favor the intermediate states M_E_ and M_I_, where the protein dwells for longer (**Fig. 2A**). There is, however, a noticeable difference between both sides of the protein, with a dwell time of 4.7 ms for the M_E_ and 11 ms for M_I_. The dwell times for the other visited states (L_E_, L_I_, H_E_, H_I_) are about equivalent (∼3 ms). The mismatch in dwell times observed at the extracellular versus intracellular reporters points to intrinsic flexibility of individual transmembrane helices, enabling faster conformational transitions at one helix end than the other.

This is further supported by the conformational landscape observed at pH 5. A two-state model on both sides of the protein now best describes the data (**Fig.2B**), On the extracellular side, it displays an interconversion between state L_E_ and state H_E_, with mean dwell times of ∼3 ms and ∼5 ms, respectively. On the intracellular side, the protein mainly dwells for about ∼15 ms in FRET state L_I_ and shortly transitions to the FRET state H_I_ for ∼4 ms. This differential occupancy of FRET states on each side of the transporter is indicative of a protein that explores a broad structural space, extending beyond simple inward–to–outward conformational transitions through rigid-body motions. Given this complex and asymmetric conformational landscape, we sought a global descriptor of the protein’s dynamics and therefore calculated the average conformational lifetime (ACL).

The average conformational lifetime (ACL) is defined as the sum of dwell times divided by the number of states. It provides a global measure of protein dynamics, reflecting the mean time required to transition between states. For the apo protein at pH 7, the average lifetime is 4.5ms, increasing to 6.9ms at pH 5. Thus, protonation of specific residues simplifies the conformational landscape (from three to two states) leading to more stable, longer-lived structural states. Altogether, our findings show that apo LmrP samples a complex conformational landscape that is asymmetric in both state populations and transition kinetics, and that this landscape is modulated by protonation.

### LmrP displays ligand-dependent conformational dynamics

Next, we investigated how the conformational landscape of LmrP reorganizes in response to antibiotics binding. We performed smFRET measurements in the presence of 2mM of antibiotics, at pH 7, followed by mpH^2^MM analyses. Four classes of antibiotics were tested: lincosamides (clindamycin), macrolides (roxithromycin), aminoglycosides (kanamycin), and β-lactams (ampicillin).

Quantitative kinetic state analysis using mpH^2^MM shows that each antibiotic presents E-S plots for both reporters distinct from the others, indicating ligand-dependent modulation of the conformational landscape. To verify that this was not an artifact due to the presence of 2 mM solute, we performed a control experiment with 2 mM glucose, which is not recognized by LmrP due to its high polarity and hydrophilicity. A three-state model was identified as the best fit on both sides of the membrane, with a conformational dynamics and kinetic profile resembling that of the apo-state (**Fig.S4**), Thus, the changes observed relative to apo are attributable to ligand binding.

Two aspects were considered for interpreting the impact of ligand binding: (i) how the conformational landscape differs from the apo condition in terms of visited states, and (ii) whether transitions between states are faster or slower on average (i.e., changes in dwell times/rate constants). First, by examining changes in the conformational landscapes, we observed that all ligands tested lead to a simplification of the conformational landscape on one side of the transporter. Indeed, while three states were sampled on each side under apo pH 7 conditions, only two states are sampled on one side and three on the other for each ligand, suggesting that ligand binding reduces the number of accessible states and the overall conformational heterogeneity.

Interestingly, the ensemble of accessible states differs as a function of the ligand. Kanamycin (**Fig. 3A**), roxithromycin (**Fig. 3C**) and ampicillin (**Fig. 3D**) data fit a two-state model on the extracellular side and a three-state model on the intracellular side, while clindamycin binding leads to the opposite profile. Notably, these differences between ligands are not reflected in their burst-averaged histograms, supporting the importance of a proper kinetic analysis. For example, the conformational landscape of clindamycin as revealed by the E-S scatter plot (**Fig. 3B**) differs markedly from that of the other ligands and apo in terms of accessible structural states. However, its burst-averaged histogram is almost identical to that of the apo, kanamycin-, and ampicillin-bound states (**Fig. 3A, D**).

**Figure 3.**
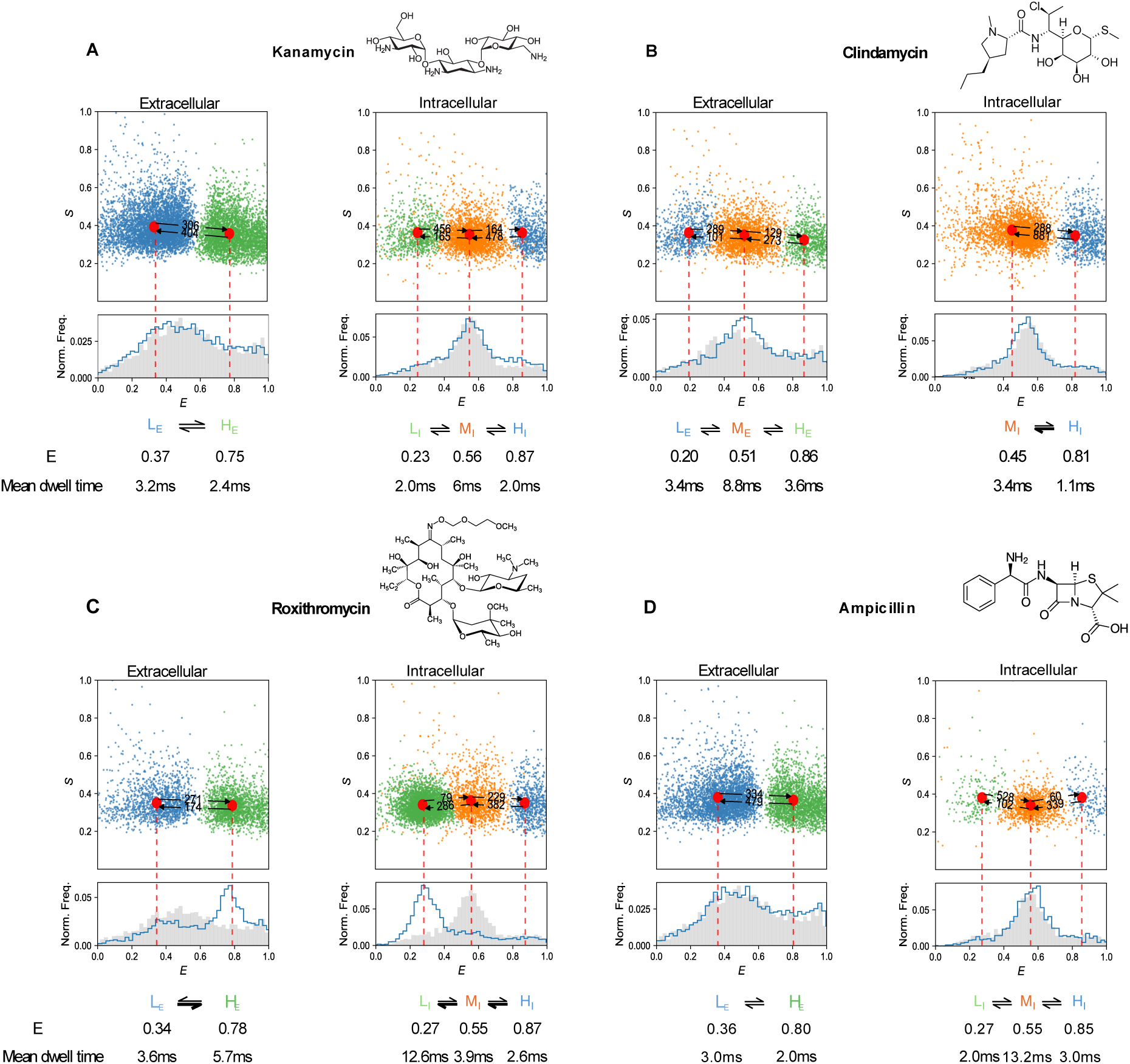
LmrP displays distinct ligand-dependent conformational dynamics. **A-D** mpH²MM analysis of LmrP in the presence of a 2 mM saturating concentration of different antibiotics. *Upper panel:* E–S scatter plots of corrected dwells, colored according to the state assignment of the selected mpH²MM model. Red circles indicate the mean FRET efficiency (E) of each state, and the numbers above the arrows denote the transition rate constants (s⁻¹) between FRET states. *Lower panel:* Burst-wise FRET histograms showing post-purified, corrected bursts. The mean E value and mean dwell time (ms) for each interconverting FRET state are indicated below. Data from the extracellular distance reporter is shown on the left, and intracellular distance reporter data on the right. The chemical structure of each antibiotic is shown adjacent to the corresponding panel (obtained from PubChem).

Analysis of kinetics of the transition between states also reveals ligand-dependent modulations, with shorter dwell times for kanamycin (ACL of 3.1 ms compared to 4.6 for apo), and longer dwell times (ACL of 5.8 ms) for roxithromycin. Clindamycin and ampicillin show ACLs (4.0 ms and 4.6 ms, respectively) close to the apo condition. Together, these data provide evidence that the conformational adaptation of this prototypical multidrug efflux pump is highly diverse and ligand dependent.

### Different kinetic profiles have different resistance profiles

We speculated that the differences between ligand-induced conformational dynamics could translate into a difference in transport efficiency, that would impact resistance. To test this, we monitored the cell growth of *L. lactis NZ9000* strain transformed with *lmrp*-containing plasmid under the control of a nisin-inducible promoter, in which the multidrug transporters LmrA and LmrCD are deleted^41,42^. The assays were done for LmrP wild-type (WT) and the transport-deficient mutant D68N, to compare the relative efficiency of antibiotic efflux as a function of cell survival at different drug concentrations^43^ ^44^.The transport-deficient D68N mutant is expressed at levels comparable to the wild type, indicating that the observed differences in relative growth arise from differences in the transport activity of LmrP^24^ (**Fig. S5**). In the presence of kanamycin (**Fig. 4A,I**), cells expressing LmrP WT required a 21-fold higher IC_50_ to inhibit growth compared to cells expressing the D68N mutant. For clindamycin, we observed 12-fold increase in IC50 for the WT, indicating a clear contribution of LmrP-mediated transport (**Fig. 4B,I**). In contrast, The IC₅₀ difference for roxithromycin was modest and did not exceed 2-folds (**Fig.4C,I**) suggesting poor transport. For ampicillin, the IC₅₀ ratio between WT and mutant strains was close to unity, indicating that ampicillin is not transported by LmrP under the tested conditions (**Fig. 4D,I**). Together, these results indicate that LmrP transports kanamycin and clindamycin, while roxithromycin or ampicillin are poorly or not transported (**Fig. 4I**).

**Figure 4.**
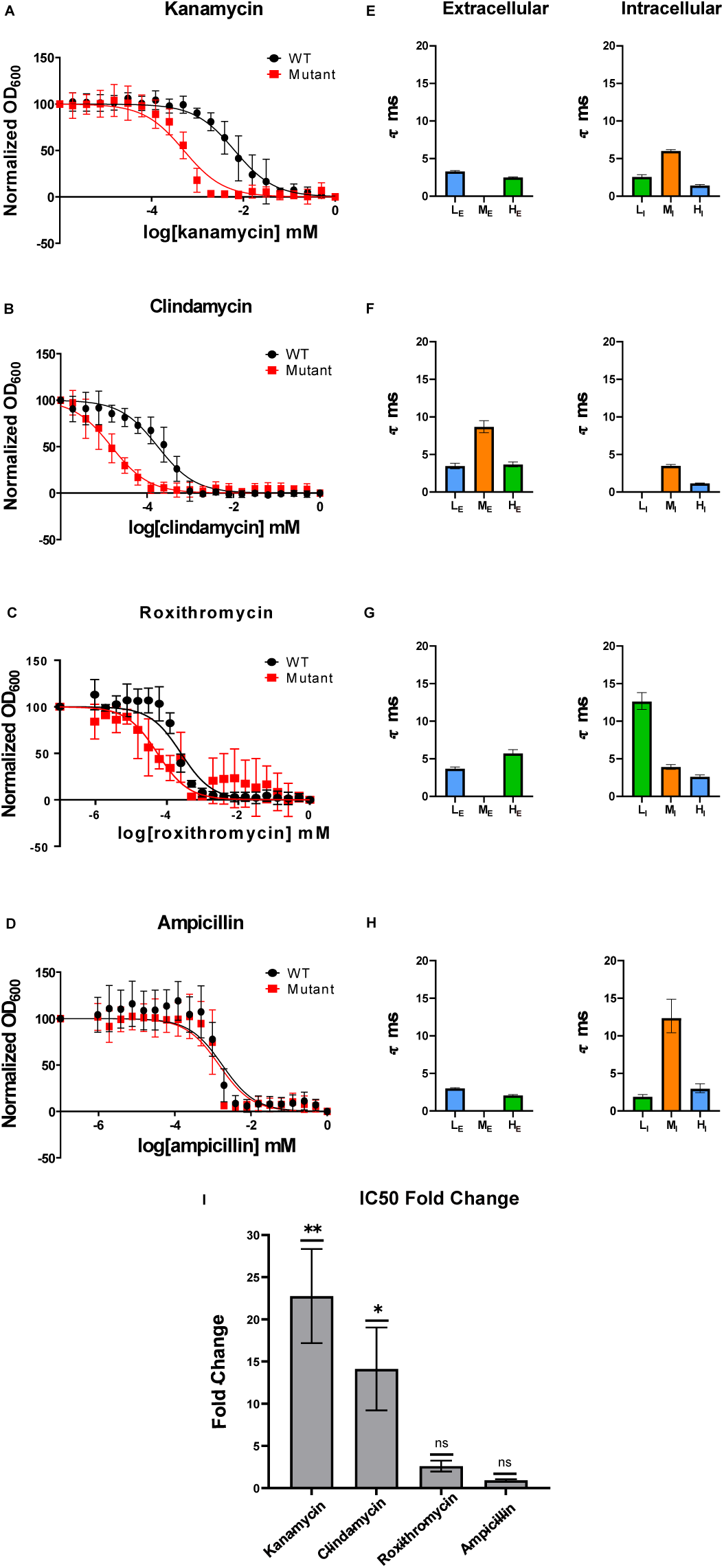
LmrP-mediated antibiotic resistance. **(A–D)** Growth inhibition assays of *L. lactis* expressing LmrP wild-type (WT, black) or the transport-deficient mutant D68N (red) in the presence of kanamycin, clindamycin, roxithromycin, and ampicillin. Non-linear regression (dose–response inhibition) was used to estimate IC₅₀ values. A representative assay is shown for each antibiotic; each data point represents the mean ± SEM from n = 4–8 technical replicates. **(E-H)**: Bar charts showing the mean dwell times in ms of different conformational states obtained from mpH²MM analysis of smFRET data in the presence of each antibiotic, for distance reporters located on the extracellular and intracellular sides of LmrP.**(I)** Bar chart summarizing the WT/Mutant fold change in IC50 values of the tested antibiotics, calculated from at least three independent biological experiments. The IC50 values were calculated using non-linear regression fit; log[inhibitor] vs response. Statistical significance was assessed using one sample t-test comparing fold changes in IC50 values of different biological replicas for WT and D68N mutant. P values are indicated as follows:*p* ≤ 0.02(**), *p* ≤ 0.05 (*), and *p* > 0.05 (ns).

We next compared antibiotic-dependent cell survival with the corresponding kinetic profiles of LmrP. Although the conformational landscape sampled by the isolated transporter in solution is unlikely to fully mirror that in cells, this comparison allows us to assess whether differences in conformational behavior are associated with functional output. Therefore, we asked two main questions: (1) Does efficient transport correlate with a specific structural sub-state(s)/ conformations? (2) Or is it rather the rate(s) of transition between states that can be correlated with transport efficiency? To facilitate this comparison, dwell times of the conformational states sampled by the extracellular and intracellular reporters were plotted alongside the cell survival curves. This representation allows visualization of which states are populated by the protein, and whether these states are long-lived, or only transiently visited. (**Fig.4E,F,G,H**).

Well-transported substrates, clindamycin and kanamycin, show different conformational landscapes both on the intracellular and extracellular reporters, with the state M_E_ visited by clindamycin-bound (**Fig.4F**) but not kanamycin-bound protein (**Fig. 4E**), while the state L_I_ is visited by kanamycin-bound (**Fig. 4E**) but not clindamycin-bound LmrP (**Fig.4F**). In contrast, kanamycin-bound protein visits the same conformational landscape (**Fig. 4E**) as the ampicillin-bound (**Fig. 4H**). This indicates that the conformations adopted or accessible upon binding do not correlate with efficient transport. In contrast, analysis of the transition kinetics between states reveals an interesting feature: the only clear difference between the conformational landscapes of kanamycin (transported) and ampicillin (not transported) lies in the faster transitions/ shorter dwell times (< 10 ms) observed on the intracellular side for kanamycin (**Fig 4E**) compared to ampicillin (**Fig. 4H**). Transition kinetics/dwell times of clindamycin-bound LmrP are also much faster on the intracellular side (**Fig. 4F**), whereas roxithromycin-bound LmrP displays long dwell times (>10ms) and slow transition kinetics (**Fig.4G**). Together, these data suggest that fast sampling of the conformational landscape on the intracellular side upon binding might be a prerequisite for efficient transport.

## Discussion

Our single-molecule FRET (smFRET) analysis shows that different antibiotics differentially reshape the conformational dynamics of the MFS multidrug transporter LmrP, giving rise to distinct ligand-dependent conformational landscapes.

LmrP and homologous multidrug transporters (e.g., MdfA, NorA, QacA) function as antiporters that exchange protons for substrates with protons entering from the extracellular medium and being released to the cytosol^45–47^, while substrates are recruited either from the inner leaflet of the membrane^48^ or directly from the cytosol^17^. Hence, the initial contact between the ligand and the transporter occurs on the intracellular side and involves different conformational adaptation for binding from different locations (inner leaflet or cytosol). Interestingly, clindamycin and roxithromycin are lipophilic, with logP values of 1.04 and 3, respectively (ChEMBL), while kanamycin and ampicillin are hydrophilic, with logP values of −7.6 and −2.0, respectively, suggesting different recruitment sites.

Remarkably, the conformational landscapes of LmrP in presence of kanamycin or ampicillin on the intracellular side are qualitatively similar to each other and to the apo condition (**Fig.2 and Fig.3**), transitioning between three states (L_I_, M_I_ and H_I_) with the M_I_ state more populated. In contrast, the conformational landscapes for the two lipophilic ligands depart markedly from the apo state; in the case of roxithromycin, the three states are present, but the L_I_ state is the most favored. For clindamycin, the L_i_ state is absent. This observation indicates that, on the intracellular side, binding of hydrophilic ligands results in minimal structural rearrangement, in contrast to lipophilic ligands, which induce significant conformational adaptation. Our work, together with that of others, points to a deviation from the canonical alternating access model of transport proposed for MFS that surmises that one conformational state is primed for binding and the other for release (see Introduction). We observe that a diversity of conformations appears compatible with substrate binding. Previous work on the multidrug transporter MdfA confirms this observation^49^. Substrate binding of the lipophilic compound TPP⁺ occurs in an occluded conformation, as shown by DEER spectroscopy while substrate binding of hydrophilic chloramphenicol was observed by X-ray crystallography in the inward-open conformation^50^.

However, binding does not necessarily result in transport. This work offers insight into the distinction between transported and non-transported ligands in a promiscuous transporter such as the MFS MDR transporter LmrP. Although the LmrP scaffold can accommodate structurally diverse substrates, efficient transport appears to require the preservation of structural flexibility on the intracellular side after binding. We correlated the kinetic profiles obtained for the isolated protein with resistance assays in bacteria and observed that higher transport efficiency was associated with a flattening of the conformational landscape charted by the intracellular reporter. Specifically, clindamycin and kanamycin, are both efficiently transported and induce accelerated transitions between states relative to the apo-state at pH 7 (**Fig. 5A, B**). This is indicated by the short mean dwell times below 6 ms for clindamycin and kanamycin (**Fig. 3A, B**), respectively, compared to up to 11 ms for the apo-state (**Fig. 2A**). On the other hand, ampicillin traps LmrP in the M_I_ state for 13.2 ms (**Fig. 3D**), while roxithromycin traps it in state L_I_ state for 12.6ms **(Fig. 3C).** Within the framework of this hypothesis, it is not the specific state that determines efficient transport, but rather the height of the kinetic barrier between states. (**Fig 5**). Further studies with additional ligands, whether transported or not, are required to confirm this hypothesis.

**Figure 5.**
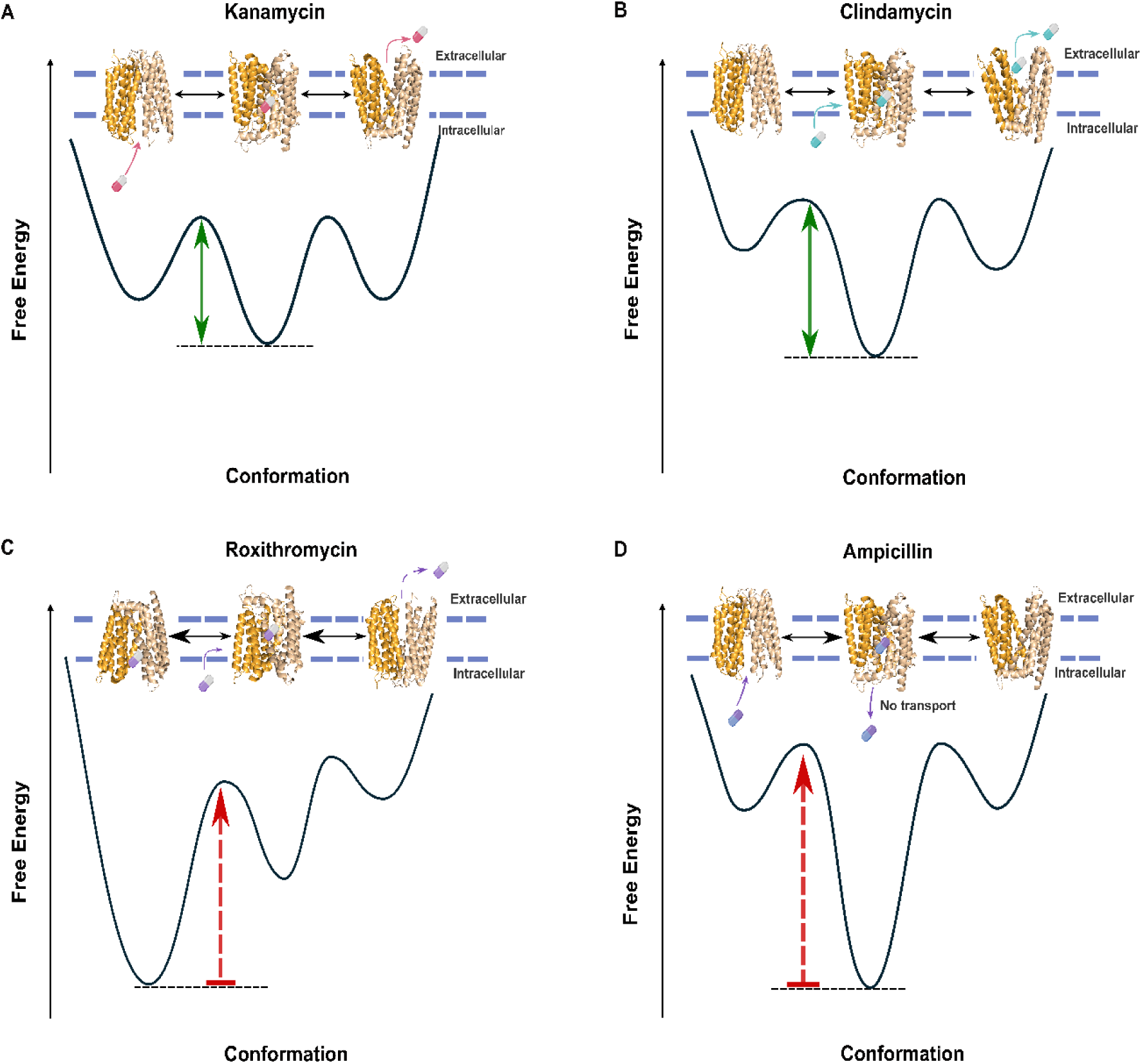
The conformational model of LmrP. **A-D** Schematic free-energy landscapes corresponding to the conformational cycle of LmrP with the antibiotics: kanamycin, clindamycin, roxithromycin, and ampicillin. The drug-bound profiles highlight distinct energetic preferences for conformational states which either lead to transport (low kinetic barrier indicated by a green arrow) or to inhibition (red kinetic barrier indicated by a red arrow). Horizontal dashed lines indicate the energy level of the most stable state in each condition. In-set is the cartoon illustration of the conformational landscapes of LmrP with each antibiotic. Cartoon illustration of LmrP conformations are based on outward-open LmrP structure (PDB:6T1Z), inward-open (published AlphaFold2 model), outward-occluded and extra-outward-open (Chai discovery model followed by manual fine-tuning, see *Methods*).

This study opens new avenues for developing strategies for functional control of multidrug resistance pumps. Our results reinforce the notion that designing ligands to target a specific conformational state (e.g., inward-or outward-facing) is unlikely to be a viable approach, as multidrug transporters can operate effectively across a broad conformational landscape. Instead, interfering with the dynamics of state interconversion i.e., kinetically trapping the transporter in whichever conformation it happens to encounter the ligand appears more promising, as stopping conformational cycling is sufficient to block transport activity. In practical terms, such kinetic trapping could be achieved by designing binders that primarily increase the activation barriers between conformational states locking moving domains, stabilizing inter-domain interfaces, thereby slowing or arresting the conformational cycle irrespective of the exact structural state that is populated. In this context, a promising proof-of-principle approach was recently demonstrated for the multidrug transporter NorA, where a Fab locked the transporter in a single conformation^43^, here the outward-open. Peptide derivatives designed from the CDR3 recognition region of this Fab efficiently inhibited NorA-mediated efflux of various drugs, including ciprofloxacin. Together, these findings, consistent with the current understanding of the remarkable plasticity of such transporters, pave the way for novel drug design strategies that move beyond a static structural view and incorporate the role of dynamic structural rearrangements.

## Materials and Methods

### Bacterial strains, plasmids and growth conditions

LmrP was expressed in the *L. lactis* NZ9000 strain that is deleted of the known ABC multidrug transporters LmrA and LmrCD^41,42^. We used the in-home pHLP5-3C plasmid, a derivative of the *L. lactis* expression vector pNZ8048 carrying the lmrP gene coding for the C-terminally 6 Histidine-tagged LmrP^51^. Briefly, cells were grown at 30 °C in M17 broth supplemented with 0.5% (w/v) glucose and 5 μg/mL chloramphenicol until the OD660 reaches 0.7. The overexpression of LmrP was then induced by the addition of 1:1000 dilution of the supernatant of the nisin-producing *L. lactis* strain NZ9700^52,53^. After 2h of induction at 30 °C, cells were harvested by centrifugation at 7000 ×g.

### Design and construction of the mutants

For each distance reporter, two residues were selected to be mutated into cysteines based on the structure of LmrP obtained by X-ray crystallography in the outward-open conformation (PDB:6T1Z) as well as on the AlphaFold predicted model of the inward- open conformation^36^. Cysteines were placed at the extracellular and intracellular extremities of chosen TMs while avoiding mutating conserved residues. The expected FRET efficiencies were estimated by simulating the fluorescent probes’ accessible volume^54^ so that they are separated by a distance within the Förster radius, 56.8 Å in this case ^55^. The accessible volume (AV) clouds were simulated using the parameters given for ATTO488 and Alexa647 in ^55^.

The plasmid template was first methylated using the dam methyltransferase (NEB), following the manufacturer’s recommendations and as described in^56^ Mutations were introduced by site-directed mutagenesis using the Pfu DNA polymerase (Promega) in the in-home pHLP5 plasmid, in which an alanine had previously replaced the only endogenous Cys270^20^. The primers were designed as described in^57^. After transformation, plasmid DNA was extracted and verified by sequencing.

### Preparation of inside-out membrane vesicles

As previously described^51^, cells were washed in 50 mM HEPES pH 7. They were resuspended (10 mL for each L of culture) in the same buffer supplemented with 7 mg/mL of lysozyme from chicken egg whites (BioChemica), 10μg/mL Deoxyribonuclease I from bovine pancreas DNAse I (VWR), 10 mM MgSO4, and 50mM Dithiothreitol (DTT). After 90min incubation at 30 °C, cells were homogenized with a Potter-Elvehjem-type tissue grinder and lysed by four passes at 1000–1500 bars using a high-pressure homogenizer. Cell debris and undisrupted cells were subsequently eliminated by centrifugation at 17,000 × g. Inside-out membrane vesicles were then isolated by ultracentrifugation of the supernatant at 100,000 ×g for 2h at 4°C. Membranes were resuspended in a pH7 buffer of 100 mM HEPES, 300 mM NaCl, 20% (w/v) glycerol (5 mL buffer/L culture) and 1mM DTT.

### Transport assays

The transport activity of LmrP mutants was assessed by measuring Hoechst fluorescence (EX 355 nm, EM 457 nm) decay over time at 30°C, based on previously published protocols^41,51^. Briefly, inside-out membrane vesicles of LmrP-expressing cells (∼1mg proteins) were diluted in transport buffer (50 mM HEPES, 2mM MgCl2, 300 mM KCl, pH 7.4) with 0.3μM of Hoechst (Invitrogen). The addition of 3 mM of ATP-Na2 in 50 mM HEPES pH 6.8 buffer supplemented with 250 mM MgSO4 allows the generation of a proton gradient by activating the endogenous F0F1-ATPase. Active extrusion of Hoechst by LmrP is detected by a decrease of fluorescence. Addition of 1μM of the ionophore nigericin dismisses the proton gradient and Hoechst is no longer actively extruded by LmrP, causing the fluorescence to increase. Raw data were normalized for fluorescence intensities relative to the initial fluorescence prior to ATP addition.

### LmrP purification and labelling for smFRET

Inside-out membrane vesicles were solubilized in a solution of 2.5% (w/V) n-dodecyl- β – D-maltoside (β-DDM) in water with 1 mM DTT. After 1.5 h incubation on a rotating wheel at 4 °C, the insoluble part was removed by ultracentrifugation at 100 000 × g for 1 h. The supernatant was batch-incubated with Ni^2+-^nitrilotriacetate affinity resin (Ni-NTA; Qiagen; 500 μL resin/L of culture) with 10mM imidazole at 4°C for 2h. The resin was pre-equilibrated with buffer A (50mM HEPES, 150mM NaCl, 10% (w/V) glycerol, 20mM imidazole, and 0.05% (w/V) β-DDM, pH 7). After the incubation, the slurry was transferred into a polypropylene column, the flow-through discarded, and the resin was washed with 8 CV of buffer A, supplemented with 0,5mM DTT. LmrP was then eluted by stepwise additions of buffer B (buffer A with 250mM imidazole and 0,5mM DTT). The more concentrated fractions were pooled together. Imidazole and DTT were removed just prior to labelling using a desalting column (Disposable PD10 desalting columns, GE Healthcare) previously equilibrated with labelling buffer (50mM HEPES, 150mM NaCl and 0.02% (w/V) β-DDM, pH 7.5) that was degassed in vacuum/N2 under continuous stirring. LmrP was concentrated to approximately 100μM using a 50kDa MWCO concentrator (Amicon, GE Healthcare) and then labelled with the donor ATTO488 (ATTO 488 maleimide, ATTO-TEC GmbH) and the acceptor Alexa647 (Alexa Fluor 647 maleimide, Life Technologies Europe BV). 250μM Alexa647 and 100μM ATTO488 were mixed in the degassed Labelling buffer and 50μM of LmrP was added for 2h at RT. Unreacted dye was removed on a desalting column, and the labelled protein was run on an SDX-200 10/300 GL (GE Healthcare) size exclusion chromatography column in SEC buffer (50 mM HEPES, pH 7, 150 mM NaCl, 10% (w/v) glycerol, and 0.02% (w/v) β-DDM). The efficiency of the labelling was checked via UV-visible absorbance at 280nm, 501nm, and 650nm to estimate the amount of LmrP, ATTO488, and Alexa647, respectively, after correcting the 280nm peak for the dyes’ absorbance.

### Confocal Multiparameter-PIE Microscope setup

The experimental measurements were recorded using a home-built multiparameter fluorescence confocal microscope operating with pulsed-interleaved excitation ^34,35,58^.

Two pulsed lasers were used for excitation with excitation wavelengths of 483nm (LDH-P-C-470, Picoquant, Berlin, Germany) and 635nm (LDH-P-C-635B, Picoquant, Berlin, Germany) with clean-up filters for both excitation sources (Chroma ET485/20x, F49-482,AHF Analysentechnik, Tübingen, Germany and Chroma z635/10x, Picoquant, Berlin, Germany). Both lasers were operated with a pulse frequency of 26.67 MHz (PDL 828 “Sepia 2”, Picoquant, Berlin, Germany) (and delayed by 18ns relative to each other. The laser beams were reflected using mirrors and dichroic mirrors into an optical fiber (60FC- 4-RGBV11-47 and PMC-400Si-2.6-NA012-3-APC-150-P, Schäfter und Kirchhof, Hamburg, Germany). After the fiber, the beams were collimated (60FC-L-4-RGBV11-4 7, Schäfter und Kirchhoff, Hamburg, Germany), passed through a linear polarizer (Codixx VIS-600-BC-W01, F22-601, AHF, Tübingen, Germany) and reflected into the backport of a microscope body (X70, Olympus Belgium NV, Berchem, Belgium) using a 3mm thick excitation polychoric (zt 405/488/561/640tpc, AHF, Tübingen, Germany) and two mirrors. The power of the excitation sources was measured and adjusted to the desired power before the beam entered the microscopy body using a laser power meter. Excitation powers of 100µW and 50µW were set for the 483nm and 635nm excitation sources, respectively. The excitation lasers were focused into the sample volume using a water-immersion objective lens (UPLSAPO-60XW, Olympus), and the confocal spot was focused about 10µm into the measured solution. The emission was collected through the objective lens, transmitted through the polychroic mirror, focused through (AC254-200-A- ML, Thorlabs, Bergkirchen, Germany) a 75 µm-pinhole (P75S, Thorlabs), and collimated again (AC254-50-A-ML, Thorlabs). Subsequently, the emission was spectrally separated using dichroic mirrors. The 483-nm emission (referred to as B) is reflected (Chroma T560lpxr, F48-559, AHF), filtered (Chroma ET525/50m, F47-525, AHF). The 635-nm emission (R) was passed through and filtered (Chroma ET705/100m, AHF). Both emission beams are furthermore separated according to their polarization (BS252, Thorlabs) and then detected using four avalanche photodiodes, two for the parallel (‖) and perpendicular (⟂) polarization components of each spectral range (PD-100-CTE, MPD for B‖ and B⟂; EG&G SPCM-AQR12/14 for R‖ and R⟂). The detector outputs were routed (HRT-82, Becker & Hickl, Berlin, Germany) to and recorded with a TCSPC card (SPC-630, Becker & Hickl). The experimental TCSPC data were recorded in a 12-bit format encoding for the total experiment time (“macrotime”), time relative to the laser excitation pulse (“nanotime”), and a detector identification signal. Data processing of the recorded TCSPC data is discussed in the section Burstwise Data Processing(^35,59^).

### Single-Molecule FRET Data Recording

Before performing sample measurements, the alignment of the smFRET microscope was confirmed by measuring a mixture of 5 nM each of ATTO488-COOH and ATTO655- COOH in water. The setup was aligned to achieve brightnesses of > 100kHz for the donor and >70kHz for the acceptor detectors with laser excitation powers of 100 and 50 µW for the donor and acceptor excitation laser, respectively. The purified and double-labeled LmrP protein was diluted in measurement buffer: 50 mM HEPES pH 7 (adjusted for 20°C, Sigma-Aldrich), 150 mM NaCl (VWR Normapur), 0.02 % β-DDM (w/V, Carl Roth) to a concentration of approximately 50 pM and to achieve a burst event frequency of 10 to 20 / s. For measurements in the presence of a substrate, the burst concentration-diluted solution of protein was allowed to incubate with the indicated substrate concentration on ice for 30 minutes. The measurement buffer was recorded without the protein to confirm the absence of an excessive and potentially interfering signal contribution. This was repeated for each substrate with the same concentration as in measurements with labeled protein. The sample solution was recorded on an 8-well Nunc Lab-Tek chambered 1.0 cover glass (Thermo Fisher Scientific). The confocal observation volume was focused about 10 µm above the surface of the cover glass into the sample solution. Measurements were recorded at room temperature (approximately 22 °C).

### Burst-wise Data Processing

The experimentally recorded TCSPC data were processed by converting the recordings to the HDF5 format using the phconvert Python module (phconvert 2024). The data were further processed using the FRETbursts module^60^. The detector identification signals and detector timing ranges, which indicate the spectral range and polarization (the photon records of the two polarization channels per spectral channels were summed) of the detected photon, were assigned to either the donor excitation/donor emission (D_ex_D_em_), donor excitation/acceptor emission (D_ex_A_em_), or acceptor excitation/acceptor emission (A_ex_A_em_) to disentangle the recorded signal according to the pulsed-interleaved excitation scheme. The background contribution was estimated using a single-exponential fit with a 30-second time window and a background threshold of less than 1.7. Individual burst events were identified using an all-photon burst search with a 20-photon sliding window and a minimum burst event acceptance threshold of 7-times the background count rate. Burst-wise parameters, such as the burst-wise FRET efficiency *(E)* or stoichiometry *(S),* were automatically calculated by the FRETbursts module (see section Calculation of Burstwise Parameters and Parameter Correction)^60^. To remove bursts that only contained signal from either the donor or acceptor dye, a minimum of 60 D_ex_D_em_ or D_ex_A_em_ photons and at least 40 A_ex_A_em_ photons were required. The distribution of bursts in *E* and *S* was assessed for the presence of a FRET-active population with *S* of around 0.5 which indicates emissions from both the donor and acceptor dyes. This removes bursts with predominantly non-emissive donor or acceptor dyes (with *S* of 1 and 0, respectively). The photon-count filtered set of bursts was loaded into the burst mpH^2^MM module for removal of short-lives donor or acceptor blinking states using a four-state mpH^2^MM fit ^61^. All photon streams (D_ex_D_em_, D_ex_A_em_, and A_ex_A_em_) were used for mpH^2^MM fitting. The fit results were used to filter out individual burst events in which a molecule ever visited a non-FRET-active state during the transition time through the observation volume. This was achieved by identifying the states with near-1 and near-0 *S* and removing such burst events from the list of bursts. This procedure removed about half of the burst events from the presented data. A second round of mpH^2^MM analysis was performed with the cleaned dataset to describe the dynamics of the data (see supplementary data for a step-by step overview of the data processing and Hidden-Markov Model Fitting)^39,60^.

### Calculation of Burst-wise Parameters and Parameter Correction

Using the pulsed interleaved excitation (PIE) scheme for smFRET, the FRET efficiency is defined as

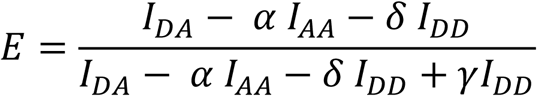

and the stoichiometry as

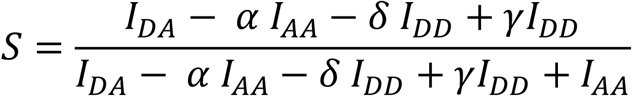

Where I_DA_, I_AA_ and I_DD_ are the background corrected number of photons detected in the acceptor detection channels after donor excitation, in the acceptor detection channels after acceptor excitation and in the donor detection channels after donor excitation, respectively; *α* is a correction factor for the direct excitation of the acceptor with the 483 nm laser; δ is a correction factor for the emission crosstalk of the donor in the acceptor channel and *γ* is a correction factor for the relative detection efficiency of the donor and acceptor ^34,35^.To account for instrument-specific difference of donor and acceptor signal detection efficiencies, the γ correction factor was determined by measuring a mixture of FRET-labeled DNA standards with two different FRET efficiencies and labeled with the same dyes as the LmrP protein (Alexa Fluor 488 and ATTO 647N). The 𝘢 and β correction values were determined from the protein measurements itself by fitting. The obtained *E-S* histogram was used to determine the corrections. The reader is referred to Maslov et al. for details on smFRET correction factors^34,59^. The correction values were found to be similar between different measurement repeats. The following correction values were used for the processing of all presented datasets: 𝘢 = 0.03, δ = 0.03, and γ = 1.2.

### Analysis of Intra-Bursts Dynamics with Hidden-Markov Model Fitting

The processed and blinking-removed set of bursts was used to assess each dataset for the presence of intra-burst dynamics. The processed and cleaned-up list of burst events (see section Burst-wise Data Processing) were loaded into the burstH2MM module (Harris et al., 2022) to quantify and characterize states visited by the FRET-labeled samples. A multiparameter H^2^MM fit was carried out (using the D_ex_D_em_, D_ex_A_em_, and A_ex_A_em_ photon streams) with a maximum of 3200 fit iterations to a maximum of five states. The integrated complete likelihood (ICL) criterion was assessed to predict the most likely number of states in the model as described by Harris et al. (2022). The model with the lowest ICL value, or where the states had a mean stoichiometry of S≈0.5, corresponding to FRET active molecules exclusively, was selected for each condition. A model was rejected once the returned mean values for S were markedly different from what can be considered a FRET-active state. This is the case, for example, if an mpH^2^MM model has split a state into two states with different *S* compared to the previous model with one state less. We rejected such models with a difference in *S* since no meaningful conformational state can be inferred from such a model. We estimated the mean dwell time for each state from the extracted transition rate constants between states. The mean state dwell time was calculated as tau_i = 1 / ((∑k_i,j) / N_states) as the inverse sum of all exchange rates that depopulate state i divided by the number of depopulation paths. The burst classifications were used to visualize the distribution based on the total burst duration (burst-wise histogram) or per-dwell basis (dwell-based scatter plot). The state-average FRET efficiency was used to compare different conditions. The uncertainty of the transition rates obtained by the best-fitting mpH^2^MM model was tested with the log-likelihood ratio test which is implemented in the burstH^2^MM module. The 95%-confidence interval was calculated using the likelihood ratio test, where the log-likelihood was reduced by 1.92 from its optimal value to determine the bounds of the interval.

We accounted for the possibility that mpH^2^MM underfitted the experimental data. However, with a simulated molecule visiting three states, underfitting with a two-state model cannot result in shorter state dwells (see Suppl. Notes). Thus, the observed increase in speed of the conformational change cannot explained by methodological artifacts but rather by an actual acceleration of the conformational landscape of the transporter.

### Cell survival assay

Mutant and WT LmrP genes were transformed into *L. lactis* strain NZ9000 ΔLmrA ΔLmrCD^42^ through electroporation. Individual colonies were grown overnight and used to inoculate 10ml cultures of M17 (supplemented with 0.5% glucose, 5 μg/ml of chloramphenicol), which were then incubated statically at 30 °C until reaching an OD660 of ∼0.7, when expression was induced through the addition of a 1:1,000 dilution of the supernatant of the nisin-producing *L. lactis* strain NZ900034^52^. After 2 h of expression, cells were diluted with M17 (supplemented with 0.5% glucose, 5 μg/ml of chloramphenicol and 1:1,000 dilution of the supernatant of the nisin-producing L. lactis strain NZ9000) to an OD660 of 0.05. These dilute solutions were then aliquoted into 96-well plates containing a dilution series of antibiotic within the M17 (supplemented with 0.5% glucose, 5 μg/ml chloramphenicol and a 1:1,000 dilution of the supernatant of the nisin-producing *L. lactis* strain NZ9000. The plates incubated to at 30 °C, without shaking, for 16 h before measuring the OD660 using Spark Multimode Microplate Reader(Tecan). The assays were conducted for at least three independent times, with six replica for each antibiotic dilution per assay. After OD normalization, the IC50 values and corresponding standard error of the mean data were calculated from a non-linear regression fit (dose–response inhibition model)^62,63^ using GraphPad Prism 10.3.

### Data availability

The code used to perform mpH2MM-based analysis of the presented experimental smFRET data is provided on Zenodo :

URL: https://zenodo.org/records/19449660

DOI: 10.5281/zenodo.19449660

(https://doi.org/10.5281/zenodo.19449660)

## Acknowledgements

This work was supported by the FNRS (MIS grant number F45322.22 and CDR J0142 24F), by an ARC Consolidator grant from the ULB (grant number 4110F000129), and the Jaumotte Demoulin Foundation for C.M. A. R. was a Research Fellow of the F.R.I.A. H.M is funded by a scientific collaborator fellowship “collaborateur scientifique” by The National Fund for Scientific Research ( Le Fonds de la Recherche Scientifique – FNRS) with grant number FC 59497. T. K. was funded by a PhD fellowship from Flemish Research Foundation (*Fonds Wetenschappelijk Onderzoek*, FWO) with grant number 11N4722N. C.G. is a Senior Research Associate of the F.R.S.-F.N.R.S.

## Author Contributions

C.G, C.M and J.H conceived the study. H.M and T.K designed the experiments. A.R performed the mutagenesis, H.M, A.R and T.K carried out the smFRET experiments and analyzed the data. C.M and H.M wrote the paper with input from all authors.

