## Supplementary Info for "Antibiotic-Specific Conformational Landscapes of a Multidrug Transporter"

### Supplementary Material

#### Supplementary Figures

**Supplementary Table.1** Expected FRET efficiencies and corresponding distances of the closed and open conformations, as predicted using FRET positioning and screening toolkit (FPS)(Kalinin et al. 2012) based on the crystal structure in the outward open conformation (PDB:6T1Z) (Debruycker et al. 2020) and the AlphaFold2 (Del Alamo, Govaerts, and Mchaourab 2021) predicted model in inward-open conformation.

| | | Outward open/Inward closed | | Outward closed/Inward open | |
|---|---|---|---|---|---|
| Residues | Localisation | $E_{expected}$ | R (Å) | $E_{expected}$ | R (Å) |
| K9C-K404C | TM1-TM12 | 0.60 | 53 | 0.30 | 65.2 |
| H98C-M374C | TM3-TM11 | 0.39 | 60.9 | 0.79 | 45.5 |

**Supplementary Table.2** Antibiotics IC50 inhibitory concentration values for *L. lactis* WT and the transport-deficient mutant D68N. The IC50 values shown are the mean values from at least three independent experiments. The WT/Mutant IC50 fold change values are also shown.

| | WT | | | Mutant | | | |
|---|---|---|---|---|---|---|---|
| | Mean IC50 µM | ±SEM | n | Mean IC50 µM | ±SEM | n | *Fold change* |
| **Kanamycin** | 16.6 | ±6.1 | 4 | 0.8 | ±0.0007 | 4 | **21.4** |
| **Clindamycin** | 0.2 | ±0.05 | 3 | 0.045 | ±3.126e-005 | 3 | **12.5** |
| **Roxithromycin** | 0.06 | ±0.07 | 7 | 0.03 | ±0.03 | 7 | **1.9** |
| **Ampicillin** | 1.8 | ±0.68 | 4 | 1.9 | ±0.4 | 4 | **0.9** |

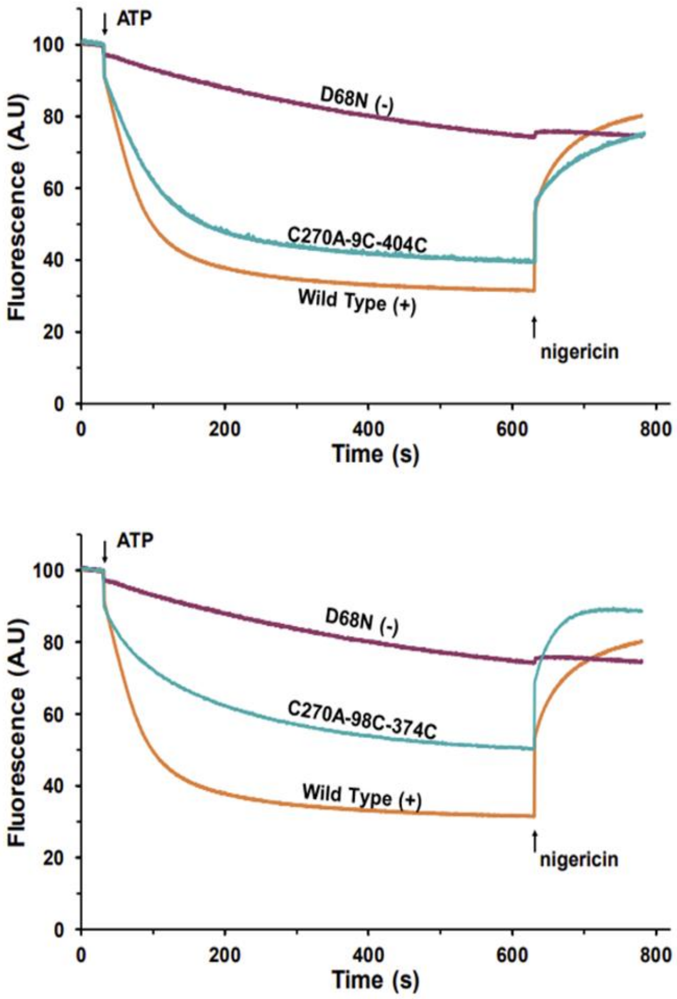


**Fig S.1**.**Transport activity assay of the double cysteine mutants.** The extrusion of Hoechst 33342 by LmrP in inside-out membrane vesicles is followed by a fluorescence-based assay (Exc: 355nm, Em: 457nm) (Steed et al. 2013). Values were normalized for fluorescence intensity. ATP energizes the endogenous F0F1-ATPase, generating a transmembrane proton gradient that activates LmrP-mediated transport. The addition of nigericin disrupts the proton gradient and inactivates LmrP transport. Hoechst is transported by LmrP WT (in orange, positive control) but not by the D68N mutant (in purple, negative control). The double cysteine mutants C270A-9C-404C (A, blue), and C270A-98C-374C (B, blue), transport Hoechst33342 as well.

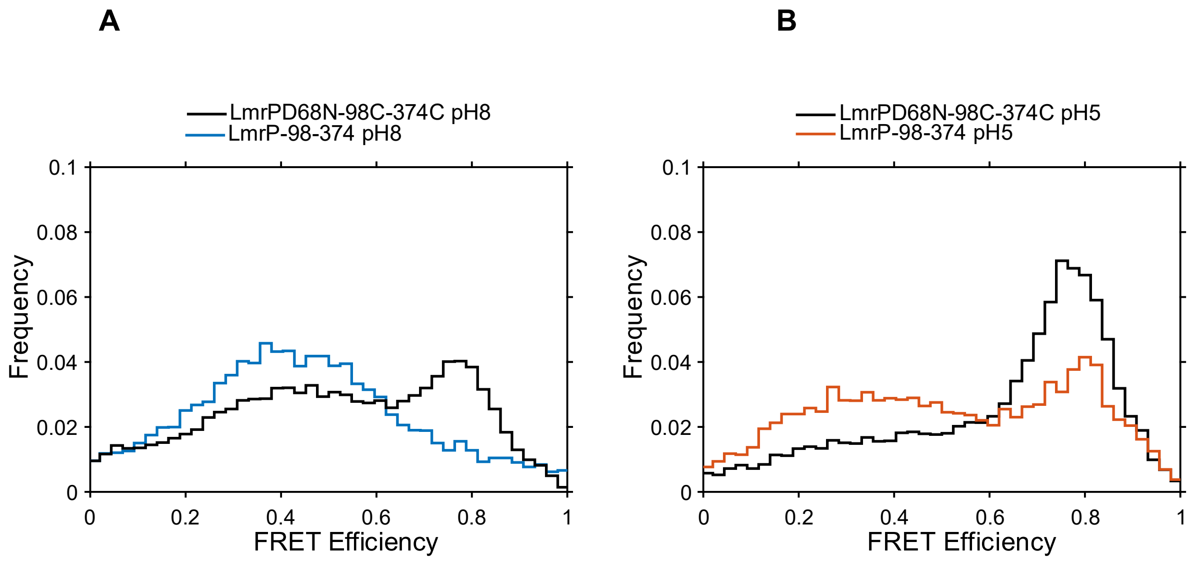


**Fig S.2. Modulation of the conformational equilibrium at the extracellular side by the D68N mutation**. The E histograms measured at (A) pH8 and (B) pH5 are shown in blue and orange respectively for the wild type, and in black for the D68N mutant .The histogram distribution for D68N mutant doesn't change with pH change like the wild type, indicating that the mutation stabilizes the protein in an inward-open conformation.

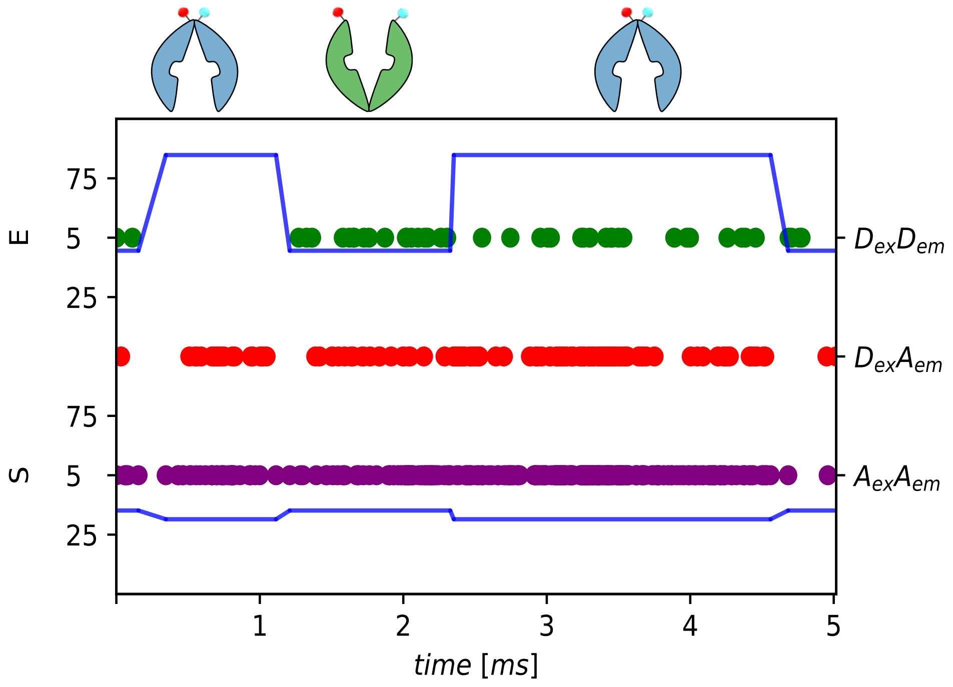


**Fig S.3. A sample photon time trace of a single burst**. Photons are represented as colored dots, with donor excitation photons colored green and red for donor and acceptor, respectively. Acceptor excitation photons are colored purple. *E* (top panel) and *S* (bottom panel) of states determined from dwells.The most likely state path predicted by the Viterbi algorithm is overlayed as a blue horizontal line. Consecutive photons with the same state are considered as a single dwell, *E* and S values are then estimated as mentioned in (Harris *et al*., 2022).

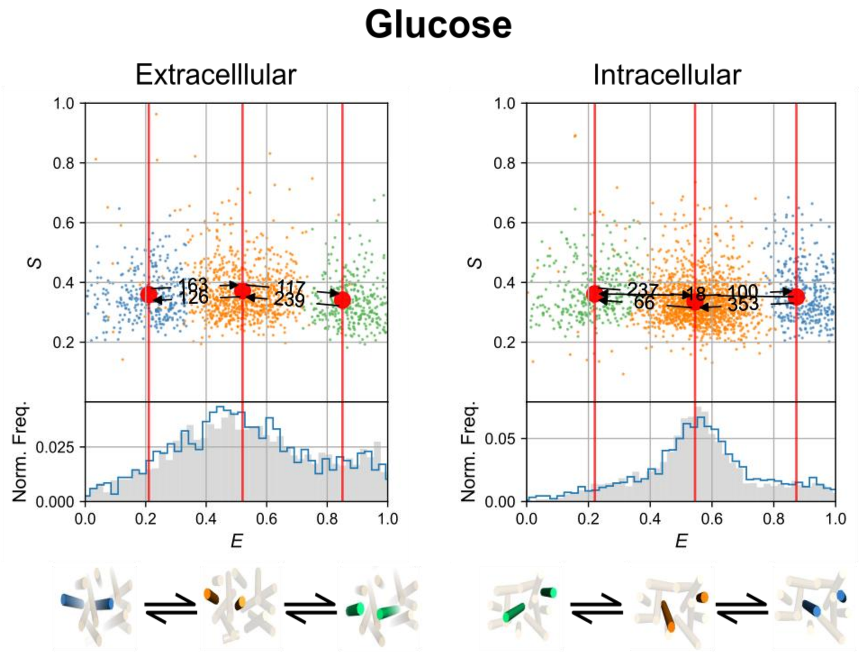


| | $L_E$ | $M_E$ | $H_E$ | $L_I$ | $M_I$ | $H_I$ |
|---|---|---|---|---|---|---|
| **E** | 0.21 | 0.55 | 0.84 | 0.22 | 0.57 | 0.88 |
| Mean dwell time | 6.3ms | 8.2ms | 4.2ms | 4.2ms | 12.5ms | 2.8ms |

**Fig S.4. Kinetic analysis of LmrP with Glucose. A.**The plots of data analysis of LmrP with glucose 2mM. Upper panel :On top the E-S scatter plot of the corrected dwells. Dwells are colored based on the assigned state of the chosen mp$H^2$MM model. Red circles represent the mean *E* value of each state, and the numbers above the arrows represent the transition rate constants ($s^{-1}$) between the two FRET states. At the bottom: Burst-wise FRET histogram showing the post-purified corrected bursts. Lower panel: Cartoon representations of the adopted confirmation for interconverting FRET states. The *E* value and mean dwell time in ms for each state are mentioned below. The extracellular distance reporter data are on the left, and the intracellular distance reporter data are on the right

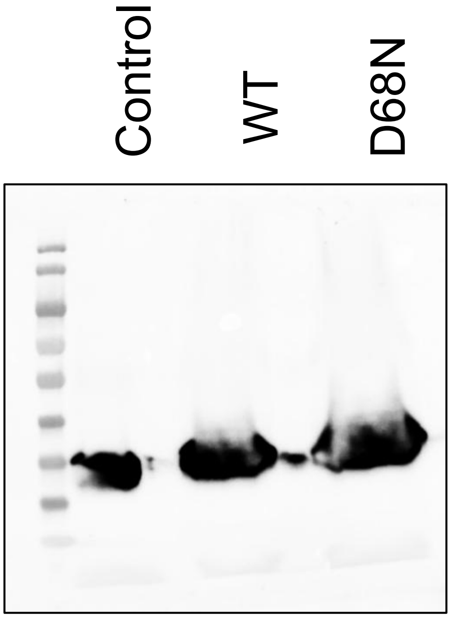


**Fig S.5.Western blot demonstrating similar levels of exp  ression for wild-type and mutant D68N of LmrP.** Resuspended membranes were prepared, as detailed in the methods. Samples of each resuspended membrane were resolved through SDS PAGE followed by Western blot analysis. Target proteins were identified using two-step, indirect detection with murine anti-His used as primary antibodies, and anti-mouse-horseradish-peroxidase (HRP) conjugates used as a secondary antibody. Antibody-bound proteins were revealed using a chemiluminescent HRP substrate.

#### Biological replica of growth inhibition assays with different antibiotics

Roxithromycin

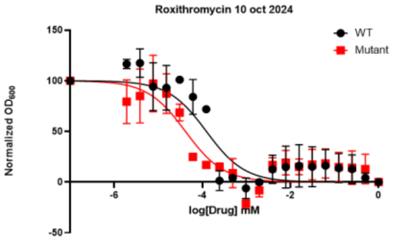


IC50 mM 0.0001234 4.129e-005

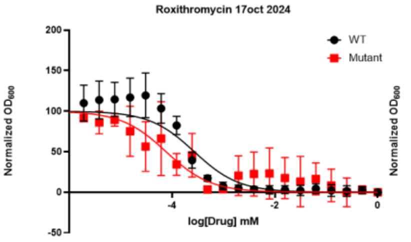


IC50 mM 0.0002696 7.615e-005

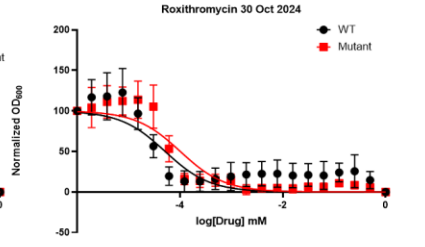


IC50 mM 5.148e-005 0.0001014

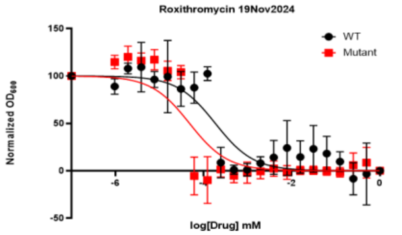


IC50 mM 0.0001838 4.773e-005

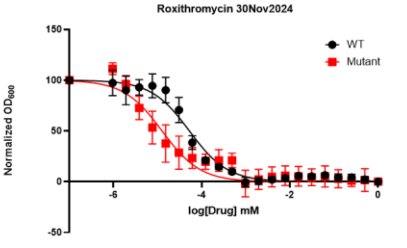


IC50 mM 5.257e-005 1.262e-005

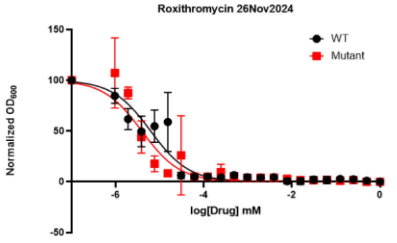


IC50 mM 6.205e-006 3.895e-006

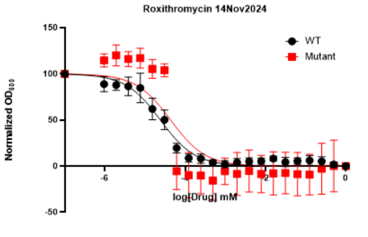


IC50 mM 2.403e-005 4.336e-005

#### Kanamycin

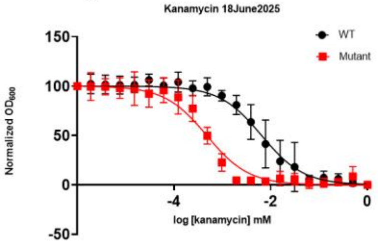


IC50 mM 0.006482 0.0004762

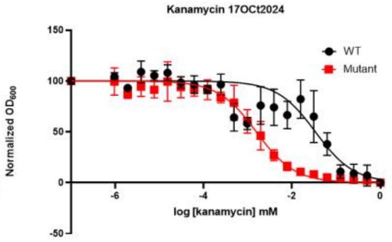


IC50 mM 0.03392 0.001556

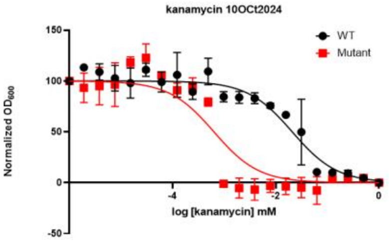


IC50 mM 0.02082 0.0006321

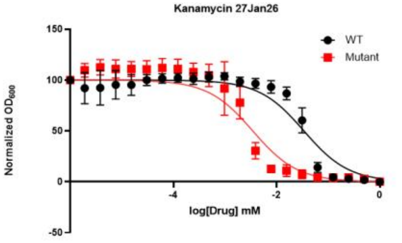


IC50 mM 0.003457 0.03392

#### Clindamycin

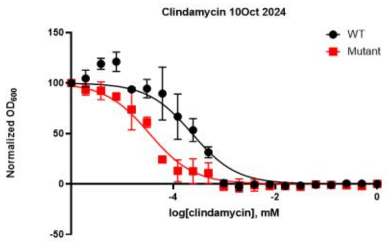


IC50 0.0002332 3.407e-005

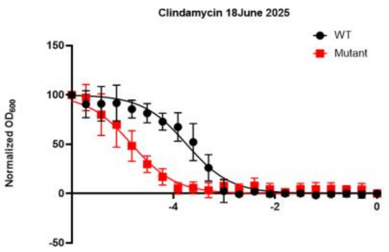


IC50 0.0002424 1.033e-005

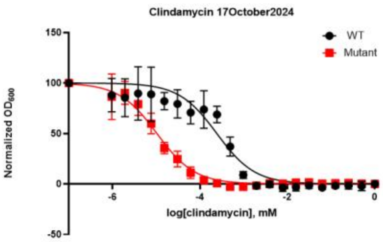


IC50 0.0001807 1.505e-005

Ampicillin

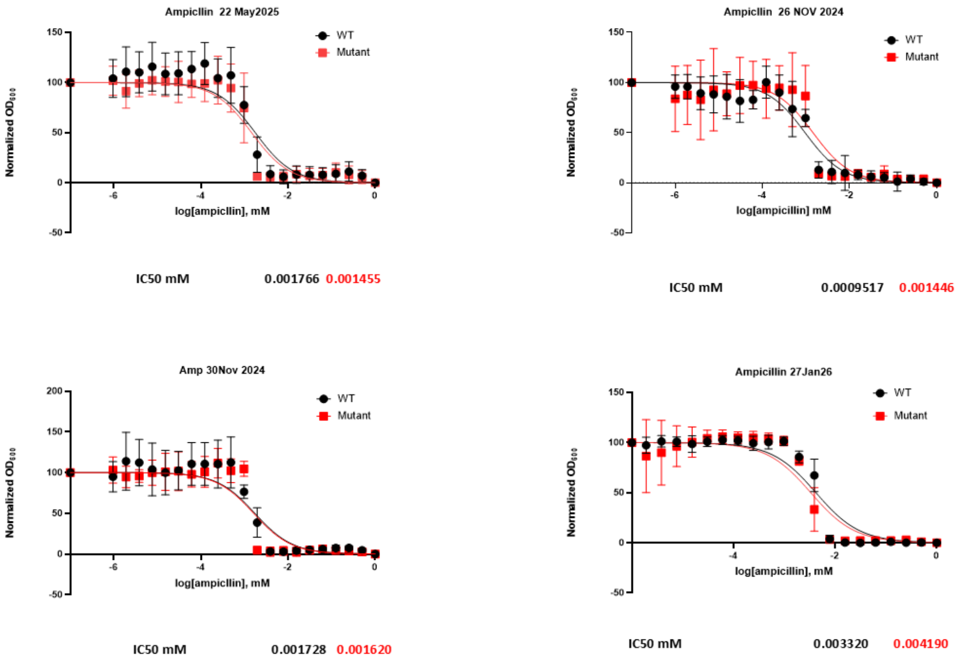


---

#### mpH2MM Analysis:

##### Determining the number of states of LmrP

The LmrP smFRET data cannot be described with standard mpH$^2$MM analysis. By that, we mean that no number of states allowed in the analysis yields an unambiguous result for the ICL and BIC' criteria. Ideally, those two criteria indicate the number of states that best describe the system. The experimental setup that was used to acquire the LmrP data was and is being used for other experimental samples for which mpH$^2$MM analysis can be done without any modification. Furthermore, we have excluded any technical origin that might explain this behavior of the LmrP data: we have acquired smFRET data of a mixture of DNA molecules that are labeled with the same dyes as the LmrP samples. For this mixture, mpH$^2$MM consistently returns a four-state model as optimal (two FRET-related states, one donor-only state, one acceptor-only state).

**Hereby, we want to showcase how the LmrP data was processed and why decisions on the exact data processing procedures were made.**

For demonstration purposes, the following experiment and analysis was chosen: 250320_LmrP_H2MM_Analysis_Template-231108_9404_pH7_apo.ipynb

Intracellular distance reporter LmrP 9-404 Apo pH7

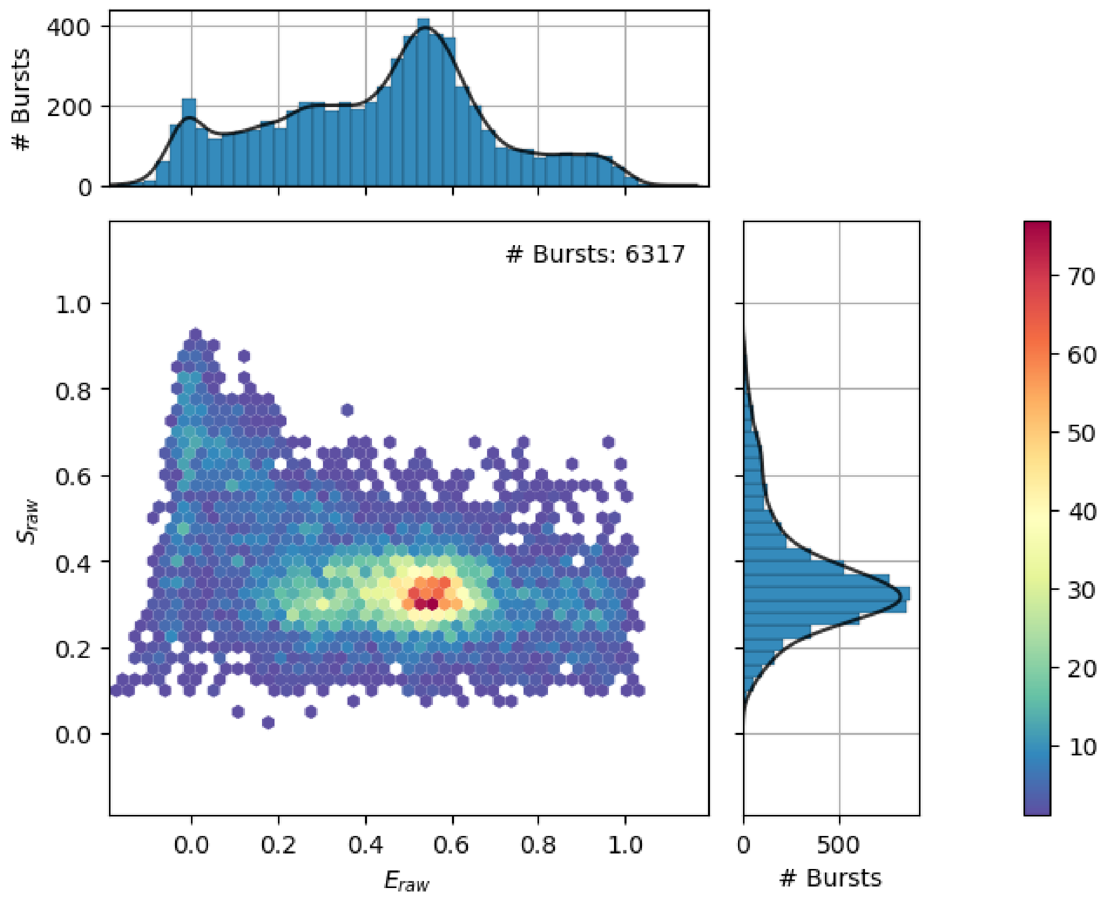


#### Standard mpH²MM analysis procedure

The processed experimental data shown above is now analyzed with mpH²MM(Harris et al. 2022). To reduce computation time, a maximum number of eight states and 2 000 fit iterations are allowed. Assuming that two blinking states are present in the dataset, up to six FRET-related states are assumed.

### ICL and BIC' plots

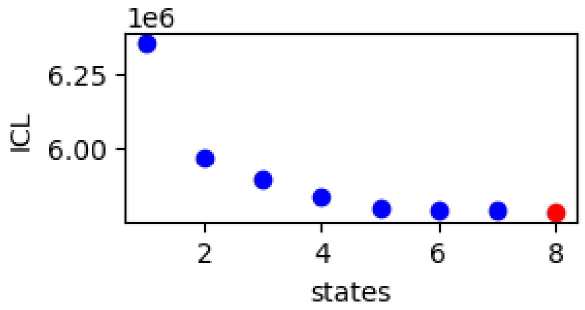

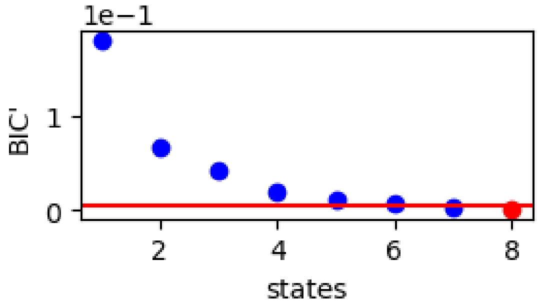


Now, we **visually** compare the dwell E-S scatterplots to find the most reasonable model.

*Three-state mode*

(1 FRET state, donor-only, acceptor-only)

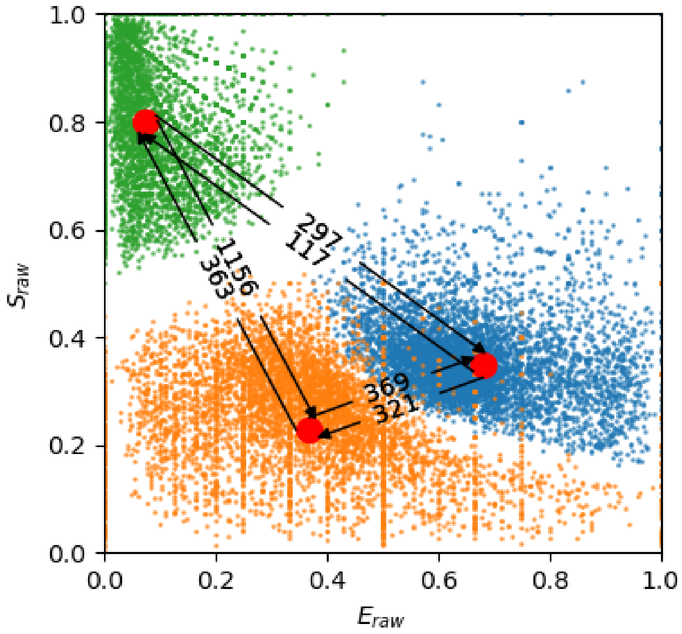


*Four-state mode*

(2 FRET states, donor-only, acceptor-only)

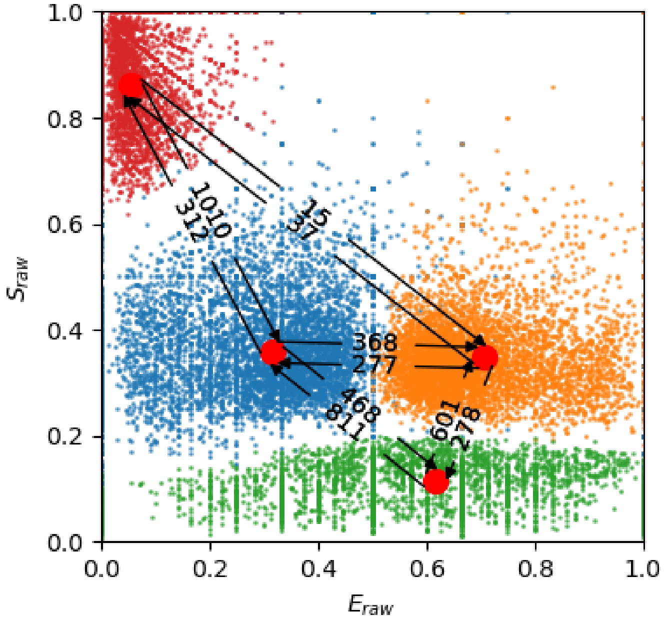


*Five-state mode*

(3 FRET state, donor-only, acceptor-only)

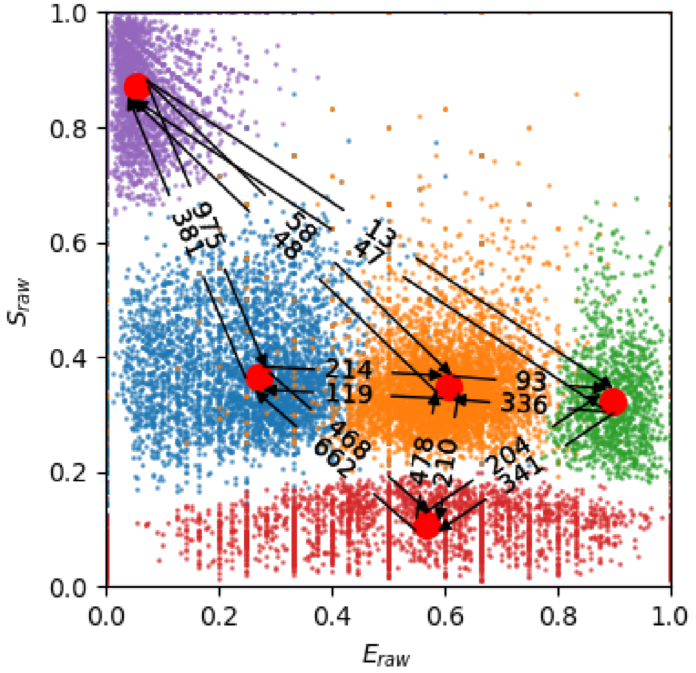


*Six-state mode*

(4 FRET state, 2 donor-only states, acceptor-only)

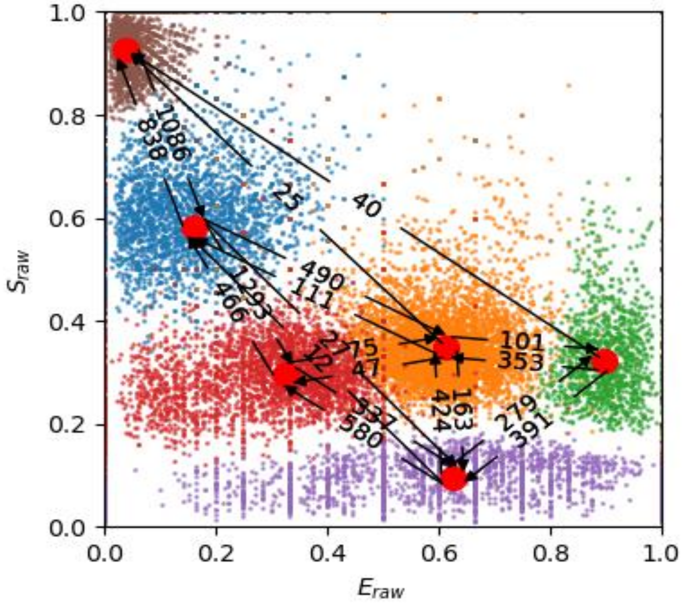


*Seven-state mode*

(4 FRET state, 2 donor-only states, 2 acceptor-only states)

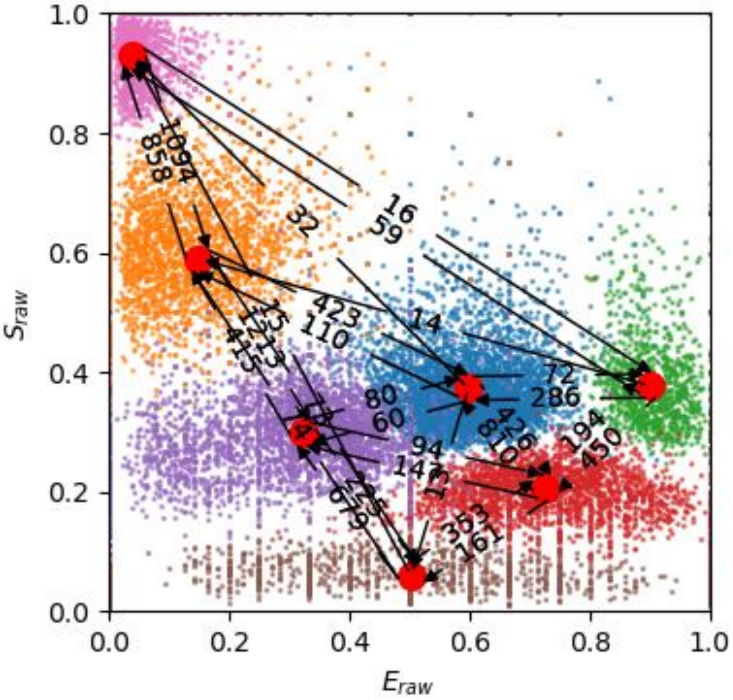


*Eight-state mode*

(4 FRET state, 2 donor-only states, 2 acceptor-only states)

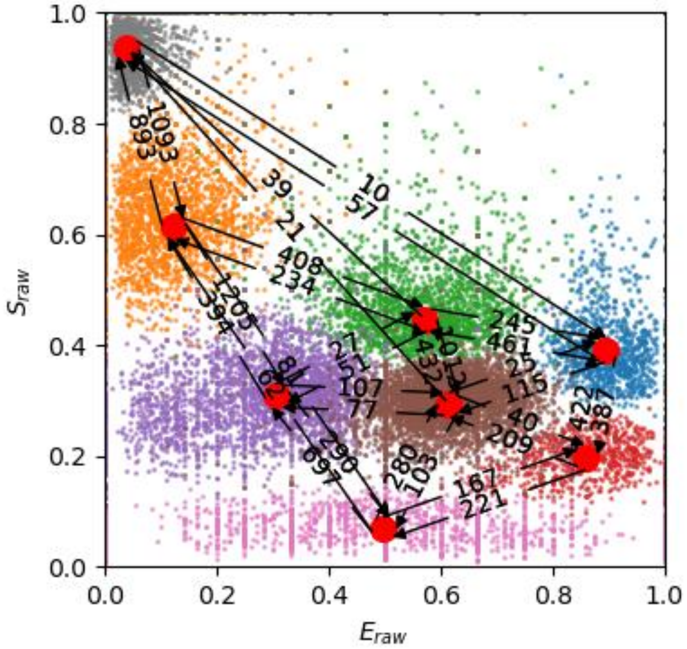


A visual comparison of the possible models shows the difficulty of choosing the correct one. If one assumes that only one donor and one acceptor state can exist, the four- and five-state models are possible. These models show a reasonable difference between blinking and FRET-relevant states along the stoichiometry axis ($S$≈0.35 for FRET-active states). Furthermore, the stoichiometries of FRET-active states appears consistent which shows that these states have similar brightnesses and thus similar emission properties of the fluorescent probes. However, there are no clear indications to favour the four-state over the three-state model. Both show reasonable transition rates much faster than what can be expected from burst frequency.

The fit obtained of the six-state model assumed that more than one donor-only state is present. The stoichiometries of the FRET-active states in this model have similar values to the states obtained from the five-state model (around $S$≈0.35). The seven-state model assumes two donor-only and two acceptor-only states and could as be reasonable as the six-state model because the FRET-active states still show similar stoichiometry values. The eight-state model can likely be ruled out because this model may be uninterpretable with some states being split purely along the stoichiometry axis (see for example the green and brown states in the eight-state model with $E$≈0.6).

#### Removal of blinking states

We attempted to remove any blinking states in the belief that standard mpH$^2$MM analysis may be interfered with by fast blinking dynamics (Kache and Hendrix 2025). We used the four-state mpH$^2$MM model from above and selected those bursts that were never identified as having transitioned into a blinking state. Assuming that the red and green states (with $S$≈1 and $S$≈0, respectively) sufficiently capture donor and acceptor blinking states, removing these bursts should leave only FRET-relevant observations, allowing to properly identify the underlying structure of the data.

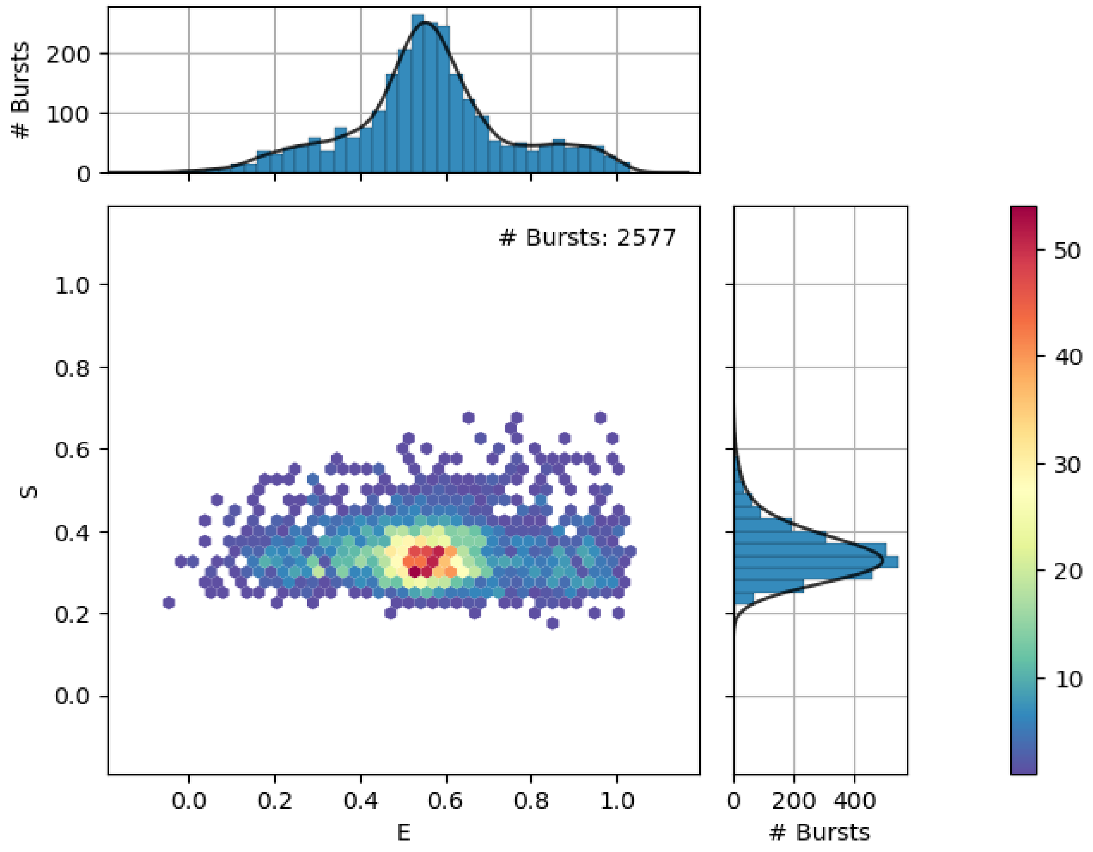


Removing blinking-affected bursts leads to a considerably decreased spread of the stoichiometry distribution (y-axis projection). Also, 2577 bursts of 6317 bursts are left after removing blinking-affected bursts, which leaves 40 % bursts. Conversely, 60 % of bursts contained blinking-related fluctuations and could possibly interfere with the ability of mpH$^2$MM to obtain the correct model from the experimental recording.

Next, we perform another round of mpH$^2$MM analysis with the cleaned dataset.

*ICL and BIC' plots*

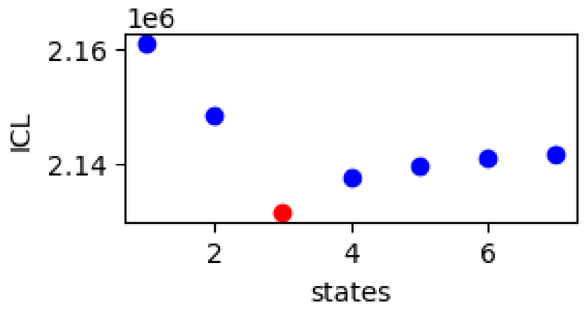

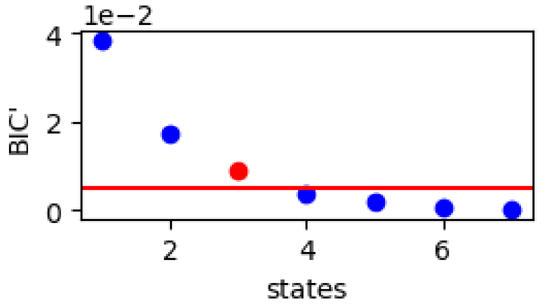


The ICL criterion clearly favors the three-state model when using the cleaned data for mpH[2]MM analysis. Assessing the two- to four-state models shows the following:

*Two-State Model*

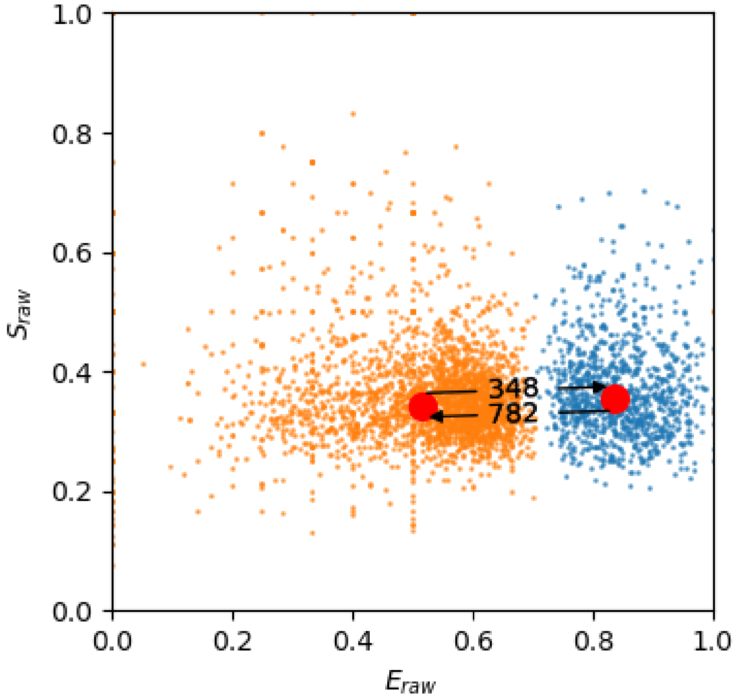

*Three-State Model*

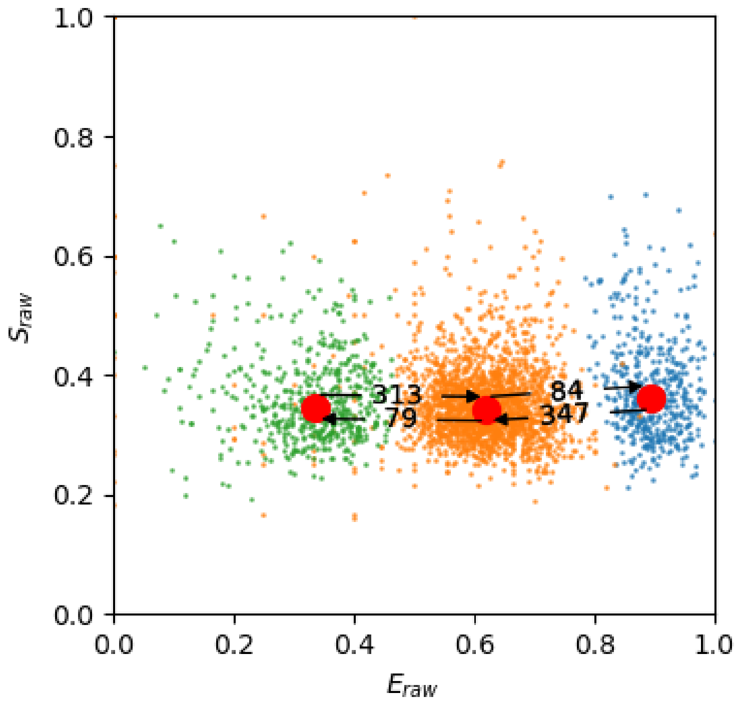


*Four-State Model*

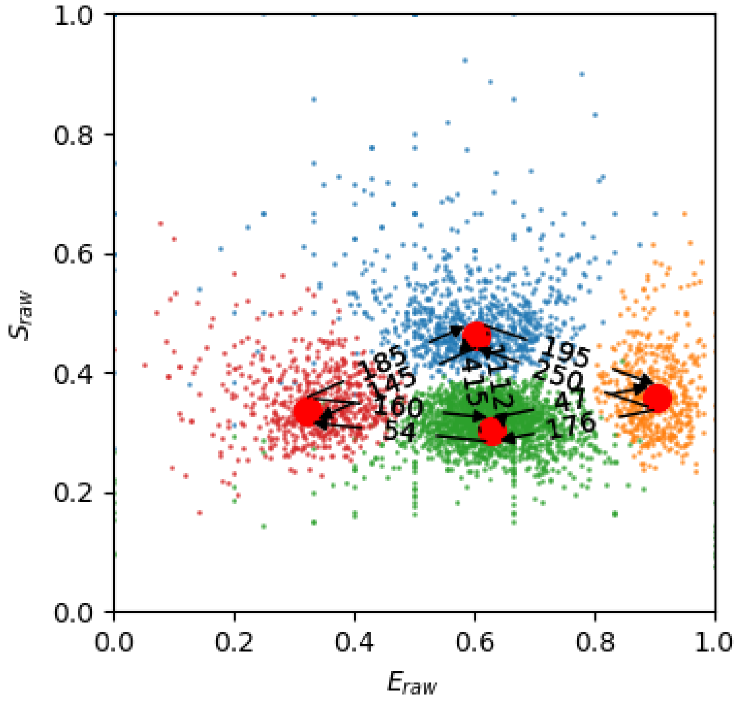


The two- and three-state models both appear reasonable with all identified states having similar stoichiometries with a value of close to 0.35. The four-state model deviates from this and starts to split the data along the stoichiometry axis. **We rule this model out**

**because we lack an explanation for this behavior and cannot interpret this in terms of an intermolecular distance between the donor and acceptor probe**.

##### Conclusion

We thus use the following criteria for the analysis of LmrP samples with smFRET and subsequent mpH$^2$MM:

1. Exclusively-acceptor and exclusively-donor bursts are removed from the dataset by requiring a minimum amount of donor-excited donor and acceptor photons and a minimum amount of acceptor-excited acceptor photons, respectively.
2. Short-lived blinking states are removed from the data by fitting a four-state mpH$^2$MM model to capture donor- and acceptor-only states. Bursts which were ever found in a blinking state are removed from the dataset and a subsequent mpH$^2$MM analysis run of **exclusively FRET-relevant** bursts is performed.
3. The ICL and BIC' criteria of the second mpH$^2$MM run are assessed. A model is ruled out, if states are **split** along the **stoichiometry axis**.

#### Underfitting a Three-State Model with mpH$^2$MM

Using a three-state model (with and without blinking), it is evaluated how underfitting with a two-state model affects the interpretation of the dataset.

We expect that the resulting exchange rates / global dynamics are being underestimated by underfitting. This is because two states are effectively merged into one and thus their dynamics are ignored by the underfitting model.

The simulations used were obtained from:

/Users/tomkache/Library/CloudStorage//Shared drives/BIOMED_DBI/TomKache/Documents/Projects/240429_H2MM_fFCS_Benchmark_Paper/data/241010_Multiparameter_Simulations/231122_TypicalBurst_Sim4/Three_State_Sims

Both files were obtained from simulated three-state exchange ($E_1 = 0.15$, $E_2 = 0.5$, and $E_3 = 0.8$) with state dwell times of 1, 2.5, and 1.5 ms, respectively. This results in a simulated exchange rate of:

- $k_{LF \to MF} = 1000 s^{-1}$
- $k_{MF \to LF} = k_{MF \to HF} = 200 s^{-1}$
- $k_{HF \to MF} = 666.6 s^{-1}$

Analysis (Ingargiola et al. 2016; Harris et al. 2022) of Simulated 3-State System without blinking

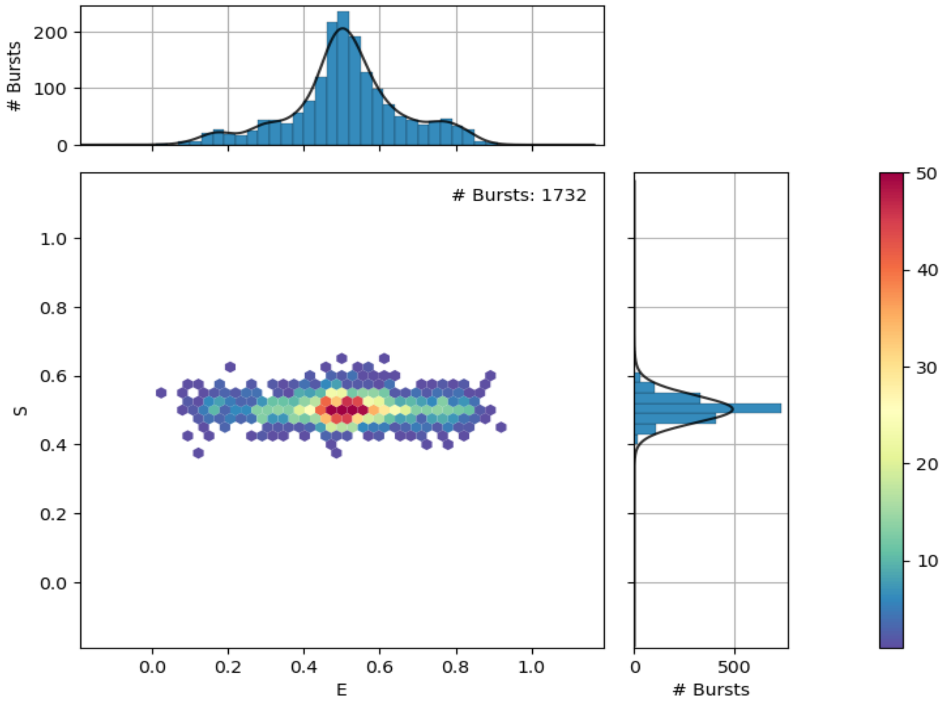


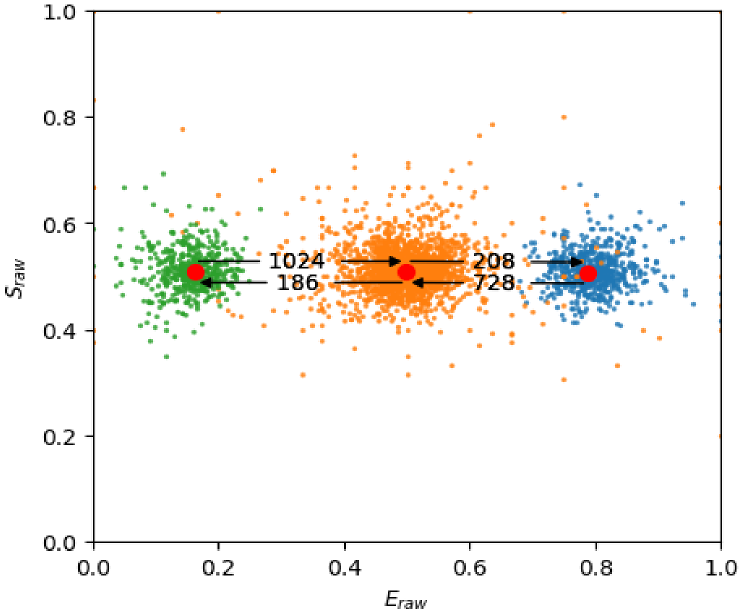


The three-state model checks out. The 'global' dynamics of the system, as described by the sum of state dwell times divided by the number of states is 1.66 ms.

Now, lets look at the two-state model:

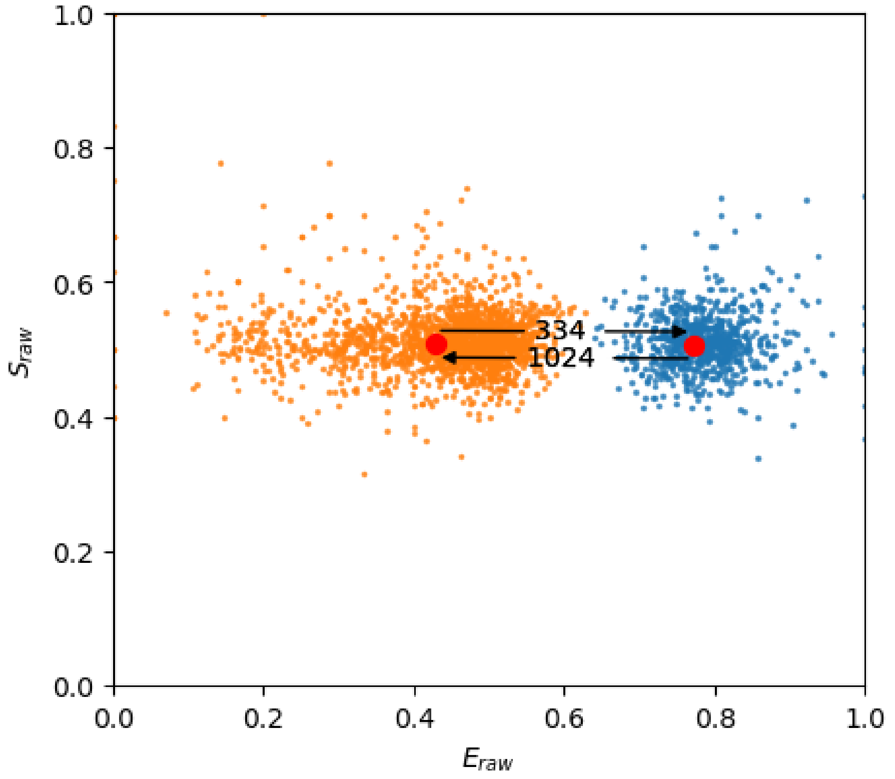


For the wrong, two-state model, we obtained a $k_{LF \to HF} = 334\ s^{-1}$ and $k_{HF \to MF} = 1024\ s^{-1}$. This corresponds to state dwell times of 2.99 and 0.97 ms. The global dynamics is therefore 1.98 ms.

The two-state model thus reports dynamics that are too slow. As hypothesized above, this makes sense, because the state transitions do not take into account a whole state but rather merged it into one single state; here the orange dwells.

#### Analysis of Simulated 3-State System with Blinking

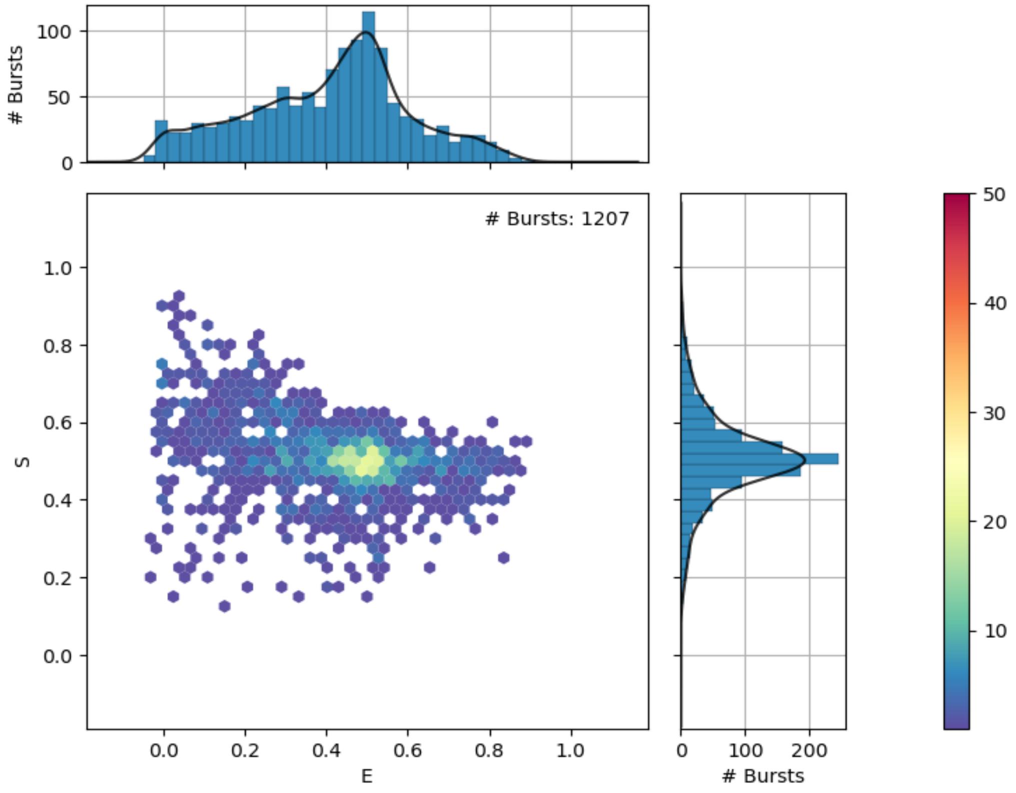


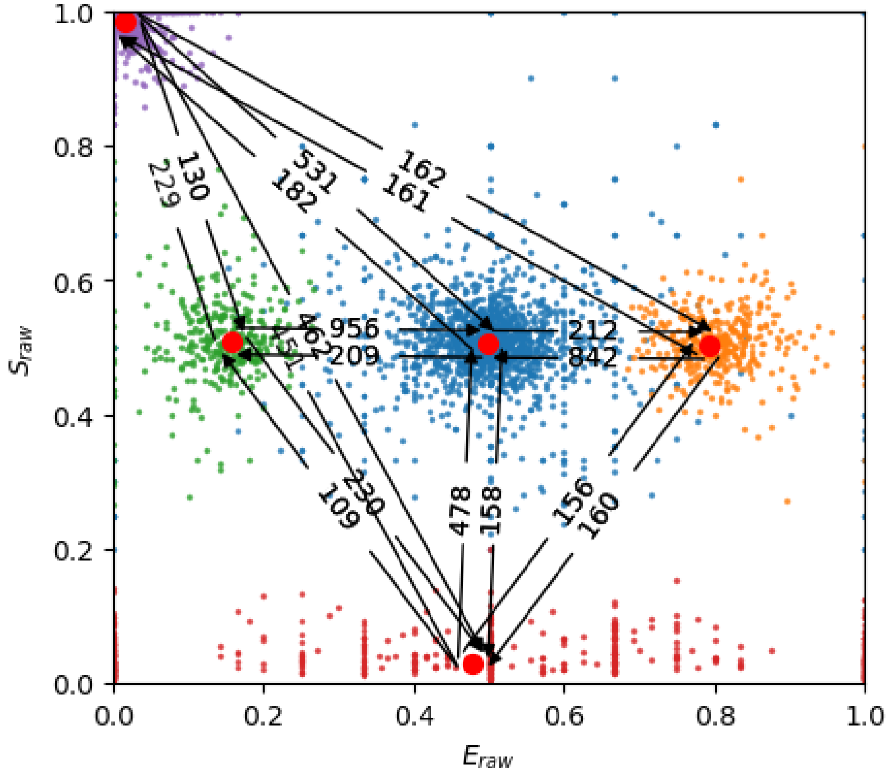

Despite the presence of blinking dynamics, the obtained exchange rates are reasonably close to the simulated values with dwell times of 1.04, 2.37, and 1.18 ms.

The underfitting model yields:

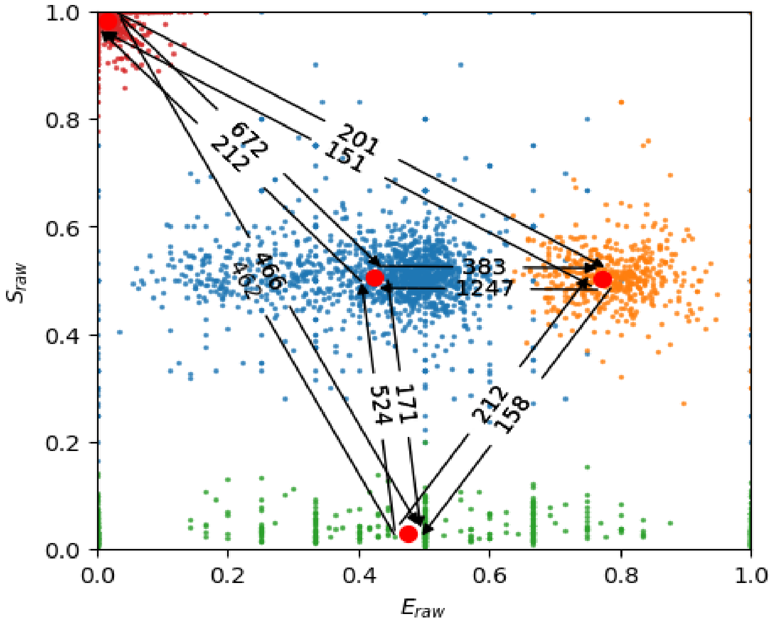


The obtained model looks similar to the simulated dataset without blinking with the LF and MF states being merged into the herein, blue state. The state dwell times are 2.6 and 0.8 ms which results in a global dynamics dwell time of 1.7 ms (compared to the actual value of 1.66 ms).

Given these simulated examples dataset without and with blinking dynamics, one can conclude that underfitting a dataset with mpH$^2$MM returns global dynamics that are too slow. This makes sense, since two states have collapsed into one state and thus lack the corresponding contribution to the global dynamics.

Conversely, we would suggest that if one observes the same sample in different conditions, and one condition yields a $N-1$ model (relative to a model of $N$ states in another condition), one can conclude that the $N-1$ model is likely correct if the global dynamics are considerably accelerated. This must be the case, since only slower global dynamics can be accounted for by having chosen the wrong model; if faster global dynamics are obtained, this is likely the results of actually having observed faster dynamics.
